# Cystatin F *(Cst7)* drives sex-dependent changes in microglia in an amyloid-driven model of Alzheimer’s Disease

**DOI:** 10.1101/2022.11.18.516922

**Authors:** Michael J. D. Daniels, Lucas Lefevre, Stefan Szymkowiak, Alice Drake, Laura McCulloch, Makis Tzioras, Jack Barrington, Owen R. Dando, Xin He, Mehreen Mohammad, Hiroki Sasaguri, Takashi Saito, Takaomi C. Saido, Tara L. Spires-Jones, Barry W. McColl

## Abstract

Microglial endolysosomal (dys)function is strongly implicated in neurodegeneration. Transcriptomic studies show that a microglial state characterised by a set of genes involved in endolysosomal function is induced in both mouse Alzheimer’s Disease (AD) models and in human AD brain and that the onset of this state is emphasized in females. *Cst7* (encoding protein Cystatin F) is among the most highly upregulated genes in these microglia. However, despite such striking and robust upregulation, the sex-specific function of *Cst7* in neurodegenerative disease is not understood. Here, we crossed *Cst7^−/−^* mice with the *App^NL-G-F^* mouse to test the role of *Cst7* in a model of amyloid-driven AD. Surprisingly, we found that *Cst7* plays a sexually dimorphic role regulating microglia in this model. In females, *Cst7*-deficient microglia had greater endolysosomal gene expression, lysosomal burden, and amyloid beta (Aβ) burden *in vivo* and were more phagocytic *in vitro*. However, in males, *Cst7*-deficient microglia were less inflammatory and had a reduction in lysosomal burden but had no change in Aβ burden. This study has important implications for AD research, confirming the functional role of a gene which is commonly upregulated in disease models, but also raising crucial questions on sexual dimorphism in neurodegenerative disease and the interplay between endolysosomal and inflammatory pathways in AD pathology.

## Introduction

Alzheimer’s disease (AD) is a chronic neurodegenerative disease characterised pathologically by buildup of protein aggregates such as amyloid beta (Aβ) and hyperphosphorylated tau. AD is also characterised by reactive glial cells, including the emergence of altered microglia. Microglia are the predominant immune cells of the brain parenchyma and play numerous roles in development and homeostasis (Colonna and Butovsky, 2017). In recent years, microglia have become heavily implicated in the pathogenesis of AD, in part due to accumulating genetic and epigenetic evidence that single nucleotide polymorphisms (SNPs) leading to altered risk for late-onset AD (LOAD) are enriched in microglial-expressed genes associated with endolysosomal function and lipid handling (Podleśny-Drabiniok, Marcora and Goate, 2020). This implies that microglia may play a causal role in the disease and supports the theory that microglia may be targeted to treat AD. In addition to genetics, sex is a major risk factor for AD, with women up to twice as likely to be diagnosed than men (Podcasy and Epperson, 2016). This sex bias is not entirely due to increased longevity in females, as increased risk remains after adjusting for lifespan, and may be due to interaction effects between sex/gender and genetics (Gamache, Yun and Chiba-Falek, 2020). The mechanisms for this, and possible interplay between microglia and sex are poorly understood.

Recently, an altered state of microglia has been identified in AD models that arises dependent on AD risk gene *TREM2*. These cells, termed disease-associated microglia (DAM)/ microglial neurodegenerative phenotype (MGnD)/activated response microglia (ARM) (Keren-Shaul *et al.*, 2017; Krasemann *et al.*, 2017; Sala Frigerio *et al.*, 2019) are characterised by downregulation of microglial ‘homeostatic’ genes *Tmem119* and *P2ry12* and upregulation of genes such as *Apoe*, *Spp1*, *Itgax*, *Clec7a* and *Cst7.* Notably, the emergence of this subset is accelerated in female mice compared to male (Kang *et al.*, 2018; Sala Frigerio *et al.*, 2019). DAM/MGnD/ARM are believed to be highly phagocytic cells due to the enrichment of phagocytic and lipid-handling pathways in their gene sets (Deczkowska *et al.*, 2018). However, the precise function of many of the genes that make up DAM/MGnD/ARM are poorly understood.

*Cst7*, coding for the protein cystatin F (CF), is amongst the most robustly upregulated genes in the DAM/MGnD/ARM signature. *Cst7*/CF expression appears to be driven by amyloid plaques as upregulation at both the RNA and protein level is spatially localized to plaques (Ofengeim *et al.*, 2017; Chen *et al.*, 2020). CF is atypical in that it is believed to play a role within the endolysosomal compartment where it acts as an endogenous inhibitor of cysteine proteases such as cathepsin L and C (Hamilton *et al.*, 2008). CF activity in neutrophils and natural killer (NK) cells has been implicated in systemic inflammation and cancer (Perišić Nanut *et al.*, 2017; Sawyer *et al.*, 2021). However, little is understood about the mechanistic role of CF in the context of central nervous system (CNS) disease. Some studies have suggested CF may play a protective role in a demyelination model by inhibiting cathepsin C (Liang *et al.*, 2016) or negatively regulating microglial phagocytosis (Kang *et al.*, 2018) but the possible disease-modifying role of *Cst7*/CF in amyloid-driven neurodegenerative disease is unknown. Additionally, although there is some evidence that *Cst7* itself is upregulated to a greater extent in female *vs.* male mice (Kang *et al.*, 2018; Sala Frigerio *et al.*, 2019; Guillot-Sestier *et al.*, 2021), the sex-specific role of *Cst7* in disease has not been investigated.

Here, we set out to investigate what, if any, disease-modifying role *Cst7*/CF may play in an amyloid-driven AD mouse model in both females and males. We first compiled data from published datasets to show that *Cst7* is indeed robustly and consistently upregulated in microglia across numerous amyloid-driven models of AD and that this upregulation appears greater in females. We then knocked out *Cst7* in male and female *App^NL-G-F^* knock-in mice which showed that *Cst7* drives sex-dependent changes in microglia at transcript, protein, and functional level. *Cst7* deletion in male mice led to downregulation of inflammatory transcripts and reduced lysosomal burden measured by LAMP2 immunostaining but had no effect on microglial Aβ burden *in vivo* or plaque deposition. However, in females, *Cst7* deletion led to increased expression of endolysosomal genes, increased lysosomal burden, and increased microglial Aβ burden *in vivo*. We further investigated this *in vitro* and showed that this is due to increased phagocytosis rather than impaired degradation. These data bring important insight to the functional role of *Cst7*, a key hallmark gene within the DAM/MGnD/ARM disease microglia signature. They also highlight that pathology-influencing effects are sex-dependent, underlining the interaction between sex and microglial function relevant to neurodegenerative disease.

## Results

### Cst7/Cystatin F is upregulated in female and male microglia in murine models of AD

*Cst7* is often detected in RNASeq or scRNASeq databases as significantly upregulated in disease models. However, an integrated analysis of *Cst7* expression in disease models stratified by sex is lacking. Therefore, we searched publicly available datasets for *Cst7* expression in microglia in mouse models of AD. We integrated data from six studies in which microglial *Cst7* expression was profiled (Fig. 1A). Expression data was restricted to microglia by use of techniques such as fluorescence-activated cell sorting (FACS) (Orre *et al.*, 2014; Wang *et al.*, 2015; Srinivasan *et al.*, 2020), immunomagnetic separation (Guillot-Sestier *et al.*, 2021), RiboTag translational profiling (Kang *et al.*, 2018) or single-cell RNA sequencing (Sala Frigerio *et al.*, 2019). Many studies did not stratify according to sex. However, we were able to collect data on *Cst7* expression in males *vs.* females in the APP/PS1 (Guillot-Sestier *et al.*, 2021) and *App^NL-G-F^* model (Sala Frigerio *et al.*, 2019). In all models, *Cst7* was dramatically upregulated in microglia in amyloid-driven disease. In models where we were able to compare sexes, *Cst7* was more highly expressed in female disease brains than in male. This is consistent with literature suggesting the transition to the DAM/ARM/MGnD microglial subset is accelerated in female mice (Sala Frigerio *et al.*, 2019). We also utilised published datasets from *in situ* sequencing (ISS), a process that allows for detection of specified transcripts with spatial resolution using barcoded probes (Chen *et al.*, 2020). Using this published dataset (www.alzmap.org) we investigated *Cst7* expression in relation to Aβ plaques in 18-month *App^NL-G-F^* mouse brain (Fig. 1B). *Cst7* transcripts (red dots) were predominantly found near Aβ plaques, a pattern mirrored by microglial-specific DAM gene *Trem2* (yellow dots) (Fig. 1). Together, these data identify *Cst7* as a key marker gene of the DAM/MGnD/ARM state and suggest that its disease-induced upregulation is related to Aβ plaque pathology.

**Figure 1:**
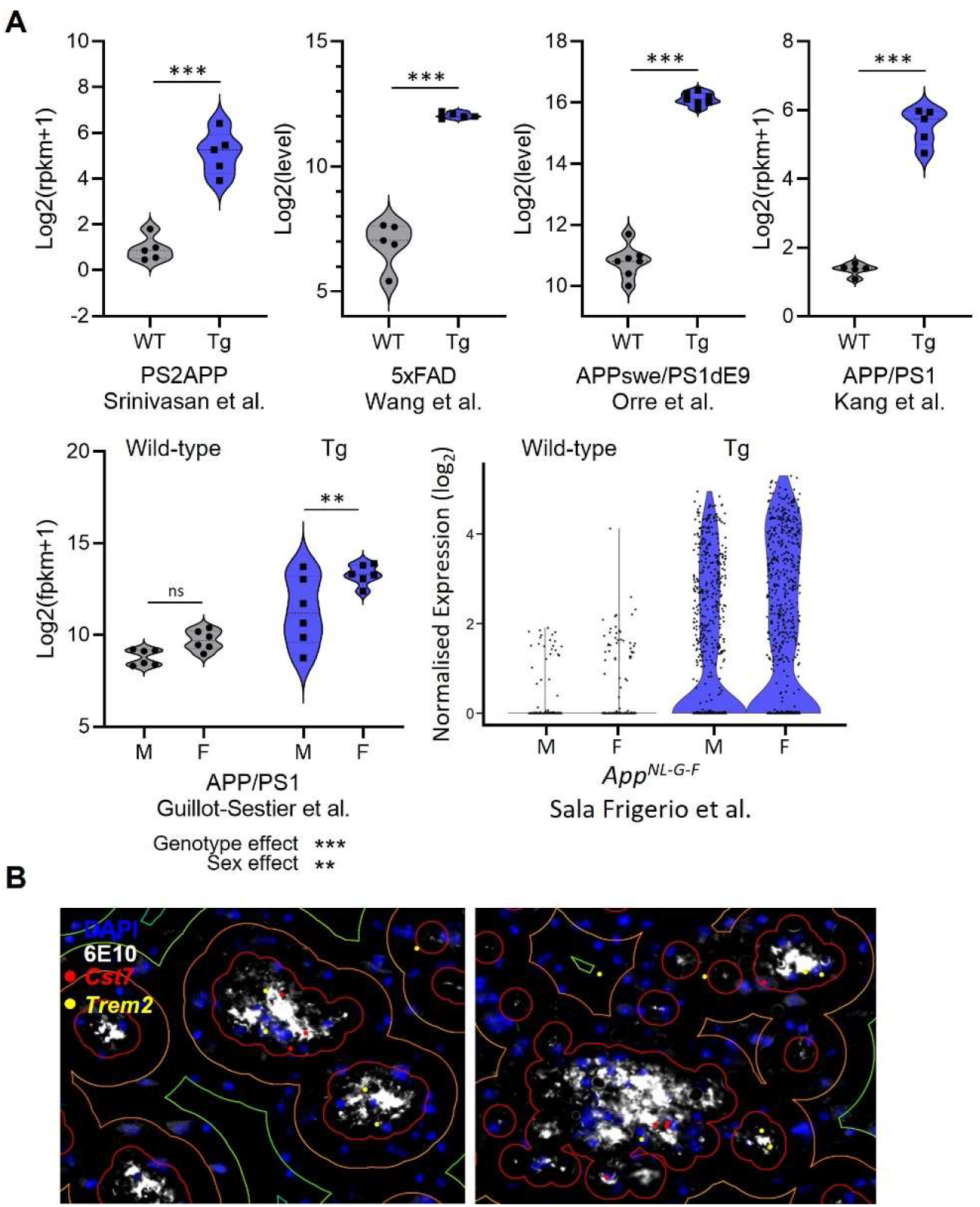
*Cst7*/Cystatin F is upregulated in microglia in murine models of AD. (A) Analysis of publicly available microglial RNASeq databases for *Cst7* expression in amyloid-driven models. Background-matched control (grey) or relevant disease model (blue). (B) Image from Chen et al. 2020 (www.alzmap.org) showing *Cst7* (red dots) and *Trem2* (yellow dots) transcripts in an 18 month old *App^NL-G-F^* mouse brain co-stained with DAPI (blue) and anti-Aβ 6E10 (white) including concentric circles around Aβ plaques. n=4-7. **p<0.01, ***p<0.001 by unpaired t-test or 2-way ANOVA with Tukey’s multiple comparisons post-hoc test.

### Cst7 deletion leads to sex-dependent transcriptomic changes in microglia in the App^NL-G-F^ model of amyloid-driven AD

As *Cst7* is so robustly upregulated in microglia in amyloid-driven disease and some studies suggest this to be greater in female mice, we next aimed to investigate whether *Cst7*/CF plays a role in regulating microglia in disease and whether this regulation could be sex-dependent. Therefore, we crossed *Cst7^−/−^* mice with the *App^NL-G-F^* model of amyloid-driven AD (Saito *et al.*, 2014) to generate background-matched non-disease (*App^Wt/Wt^Cst7^+/+^*), disease (*App^NL-G-F^Cst7^+/+^*) and disease knockout (*App^NL-G-F^Cst7^−/−^*) mice. As *Cst7* expression is limited to the myeloid compartment in the brain (Zhang *et al.*, 2014), this study primarily tests the role of microglial/border-associated macrophages in amyloid-driven disease. We first investigated the role of *Cst7*/CF in regulating the microglial response in disease by unbiased analysis of the microglial transcriptome. Mice were aged to 12 months to drive Aβ plaque burden and a DAM/ARM microglial profile that includes marked upregulation of *Cst7* before microglia were isolated from hemi-brains by fluorescence-activated cell sorting (FACS) (Fig. 2A). Microglia were defined by FSC/SSC profile, singlet, live, CD11b+, Ly6C-, CD45+ (Fig. S1) and sorted along with phenotypic markers P2Y12, MHCII and CD11c (Fig. S1) before RNA purification, sequencing and analysis using DESeq2 (Love, Huber and Anders, 2014).

**Figure 2.**
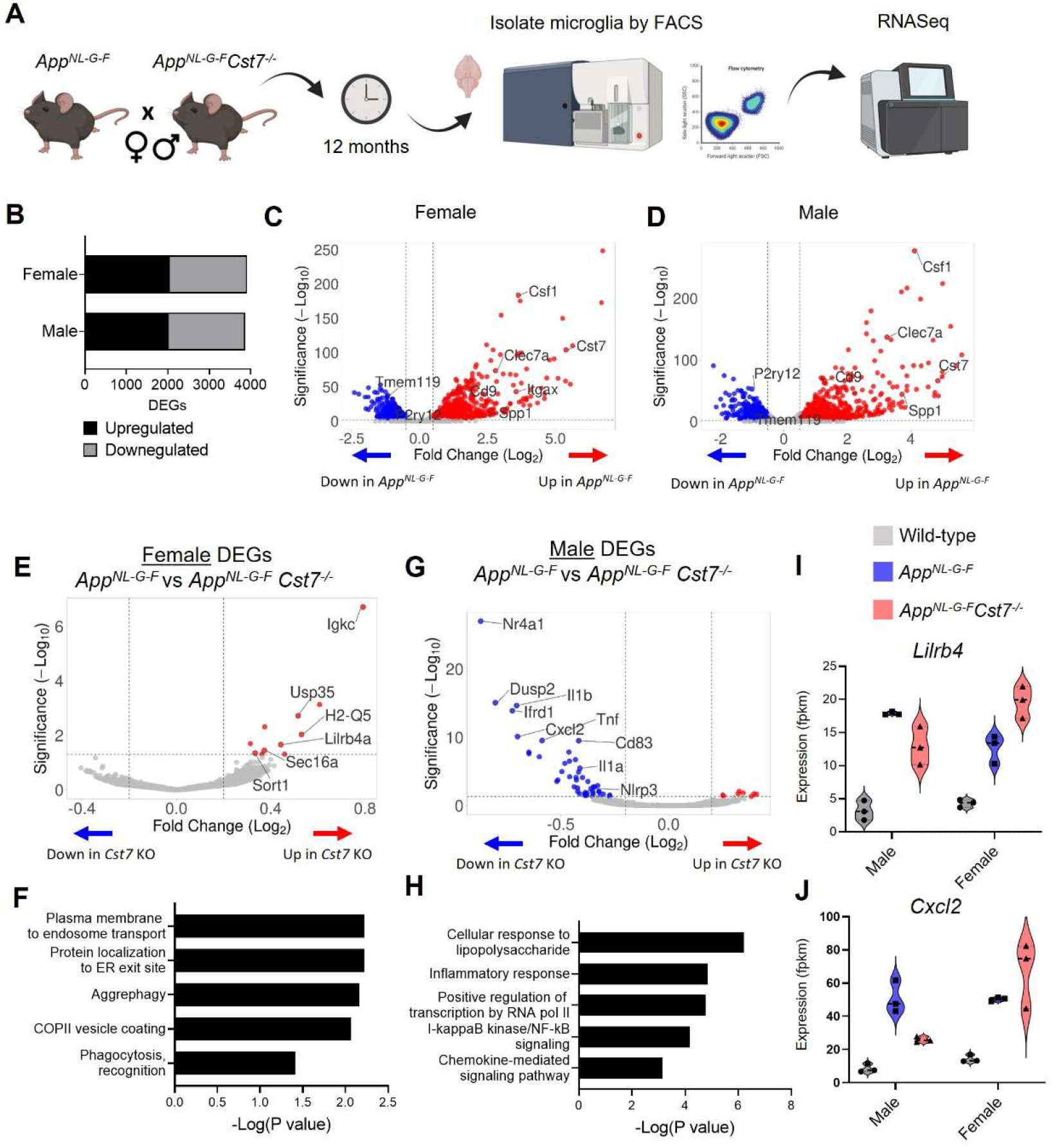
*Cst7* knockout leads to sex-dependent transcriptomic changes in microglia in the *App^NL-G-F^* model of amyloid-driven AD. (A) Study design schematic. (B) Differentially expressed genes (DEGs) between wild-type and *App^NL-G-F^* microglia in male and female mice. (C) Volcano plot of RNASeq from wild-type *vs. App^NL-G-F^* female microglia. Points in red are significantly upregulated in disease. Points in blue are significantly downregulated. Selected genes are annotated. (D) Volcano plot of RNASeq from wild-type *vs. App^NL-G-F^Cst7*^+/+^ male microglia. (E) Volcano plot of RNASeq from female *App^NL-G-F^Cst7*^+/+^ *vs. App^NL-G-F^Cst7*^−/−^ microglia. Points in red are significantly upregulated in *App^NL-G-F^Cst7*^−/−^. Points in blue are significantly downregulated. Selected genes are annotated. (F) Selected GO:BP terms that are significantly enriched in the upregulated genes from (E) and corresponding significance p value. (G) Volcano plot of RNASeq from male *App^NL-G-F^Cst7^+/+^ vs. App^NL-G-F^Cst7*^−/−^ microglia. (H) Selected GO:BP terms that are significantly enriched in the upregulated genes from (G) and corresponding significance p value. (I&J) Example gene expression (fpkm) from RNASeq of microglia isolated from male and female wild-type, App^NL-G-F^*Cst7^+/+^* and App^NL-G-F^*Cst7*^−/−^ mice. Selected genes are *Lilrb4* (I) and *Cxcl2* (J). n=5-12 mice per group, samples were prepared as 3 pools of 1-4 mice for RNASeq.

We first assessed disease-induced microglial changes by comparing *App^Wt/Wt^Cst7^+/+^* and *App^NL-G-F^Cst7^+/+^* microglia in male and female mice. *App^NL-G-F^* knock-in led to numerous changes in gene transcription with 2022 genes upregulated and 1849 downregulated in males and 2053 upregulated and 1857 downregulated in females (Fig. 2B). As expected, upregulated genes included DAM/ARM genes such as *Clec7a*, *Spp1*, *Itgax* and *Csf1,* whilst downregulated genes included microglial ‘homeostatic’ genes *P2ry12*, *Tmem119* and *Gpr34* in both sexes (Fig. 2C&D). *Cst7* was among the top ten most highly upregulated genes in males and females. Interestingly, although there were relatively few differentially expressed genes (DEGs) between male and female mice in the *App^NL-G-F^Cst7^+/+^* group (Table S1A), these genes did include DAM/ARM genes such as *Gpnmb*, *Spp1* and *Ctse*. This is consistent with the observation that the DAM/ARM profile is accelerated in females compared to males (Sala Frigerio *et al.*, 2019).

Having confirmed the upregulation of the *Cst7* and the DAM/ARM profile in this model, we next tested the role of *Cst7* in disease by comparing differential expression between the *App^NL-G-F^Cst7^+/+^* and *App^NL-G-F^Cst7^−/−^* groups in both sexes. Importantly, *Cst7* itself was the most differentially expressed gene in both sexes, confirming successful deletion (Fig. S2A). *Cst7* deletion led to significant changes in microglial transcriptome. In females, *Cst7* knockout led to upregulation of 15 genes and downregulation of two genes (including *Cst7*) (Fig. 2E). Despite the modest number of differentially expressed genes, we observed that many of the upregulated genes appeared to be involved in endolysosomal-related processes such as protein trafficking to organelles (*Sort1*, *Sec16a*) and antigen-presentation (*Lilrb4*, *Igkc*). Enrichment analyses revealed that female *App^NL-G-F^Cst7^−/−^* microglia were significantly enriched in GO:BP (Gene ontology: biological process) terms such as ‘plasma membrane to endosome transport’, ‘aggrephagy’ and ‘phagocytosis’ against their *App^NL-G-F^Cst7^+/+^* counterparts (Fig. 2F).

Next, we tested whether *Cst7* knockout led to the same effect in males as in females. Surprisingly, we observed that male *App^NL-G-F^Cst7^−/−^* microglia exhibited markedly different transcriptome compared to *App^NL-G-F^* with upregulation of eight genes and downregulation of 51 genes (including *Cst7*) (Fig. 2G). In contrast to females, downregulated genes in males were dominated by classical proinflammatory mediators such as *Il1b*, *Il1a*, *Tnf* and *Cxcl2* and enrichment analyses revealed downregulation of GO:BP terms such as ‘cellular response to lipopolysaccharide’, inflammatory response’, and ‘NF-kappaB signaling’ (Fig. 2H). Notably, many disease-induced genes followed an intriguing pattern of being suppressed in *App^NL-G-F^Cst7^−/−^ vs. App^NL-G-F^Cst7^+/+^* in males but potentiated in *App^NL-G-F^Cst7^−/−^ vs. App^NL-G-F^Cst7^+/+^* in females. Examples of this are immunoglobulin receptor gene *Lilrb4a* and inflammatory gene *Cxcl2* (Fig. 2I&J). Interestingly, *Cst7* also appeared to drive a sex-genotype interaction effect. We determined male *vs.* female differentially expressed genes in *App^NL-G-F^Cst7^+/+^* and their *App^NL-G-F^Cst7^−/−^* counterparts. In *App^NL-G-F^Cst7^+/+^* microglia, there were 33 DEGs between male *vs.* female approximately evenly split between transcripts enriched in males *vs.* those enriched in females, whereas in *App^NL-G-F^Cst7^−/−^* we observed 240 DEGs dominated by those enriched in females. These transcripts included inflammatory genes such as *Il1b*, *Tnf* and *Cxcl2* (Fig. S2B).

Together, these data show that *Cst7* drives sex-dependent effects on microglial transcriptome in an amyloid-driven mouse model of AD. *Cst7* knockout led to an upregulation of a small set of endolysosomal-enriched genes in female microglia but a striking downregulation of inflammatory genes in male microglia. The number of DEGs between *App^NL-G-F^*males and females was markedly greater in the *Cst7^−/−^* state, implying a sex-genotype interaction effect i.e. the impact of *Cst7* deficiency on microglial phenotype in *App^NL-G-^*^F^-driven disease differs quantitatively and qualitatively (e.g. in endolysosomal and inflammatory pathways) according to sex.

### Cst7 deletion leads to region and sex-dependent changes in lysosomes in App^NL-G-F^ mice

Following our investigation into the *Cst7*-dependent effects on microglia at the transcriptome level, we next sought to study changes at the protein level with spatial resolution. This allowed us to investigate brain region-dependent effects. We took sagittal brain sections from 12-month male and female *App^Wt/Wt^Cst7^+/+^*, *App^NL-G-F^Cst7^+/+^* and *App^NL-G-F^Cst7^−/−^* mice and immunostained for IBA1 to visualise microglia, anti- Aβ clone 6E10 to visualise Aβ, and, as both previous literature and our RNASeq data in Fig. 2 implicated lysosomes in *Cst7* function, LAMP2 to visualise lysosomes. We assessed three brain regions: cortex, whole hippocampus, and the dorsal subiculum (Fig. S3A), to investigate the effects of disease and *Cst7* knockout on microglial burden and lysosomal activation. As expected, knock-in of the *App^NL-G-F^* construct led to marked build-up of 6E10+ Aβ plaques, an increase in IBA1+ microglial burden around these plaques, and upregulation of LAMP2 in microglia predominantly in plaque-associated microglia in both males and females (Fig. 3A). LAMP2 predominantly colocalised with IBA1, suggesting that expression is predominantly in mononuclear phagocytes rather than neuronal cells or macroglia. Next, we compared *App^NL-G-F^Cst7^+/+^ vs. App^NL-G-F^Cst7^−/−^* male and female mice. We observed a trend towards increased IBA1+ microglial burden in *App^NL-G-F^Cst7^−/−^* mice in all brain regions, statistically significant in the subiculum (Fig. 3B-D), more evidently in male mice. Importantly, this was not only due to an increase in Aβ plaque load (which correlated with microglial burden) as images were quantified in areas of even plaque coverage (Fig. S3B-D).

**Figure 3.**
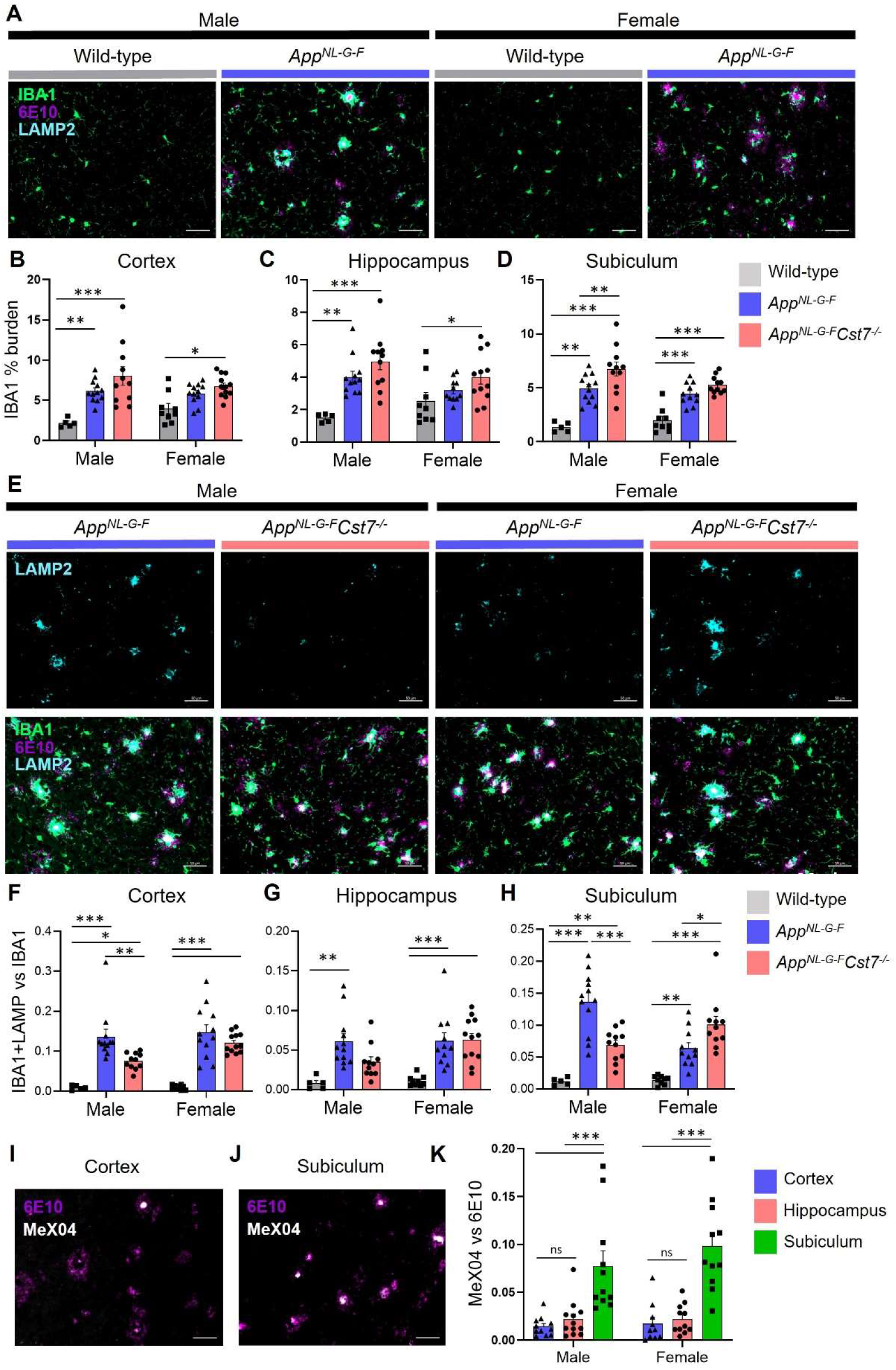
*Cst7* knockout leads to region and sex-dependent changes in lysosomes in App^NL-G-F^ mice. (A) Example images from male and female wild-type and *App^NL-G-F^* cortex stained with IBA1 (green), 6E10 (magenta) and LAMP2 (cyan). (B-D) IBA1 % coverage from images taken of wild-type, *App^NL-G-F^Cst7^+/+^* and *App^NL-G-F^Cst7*^−/−^ brains in cortex (B), hippocampus (C), and dorsal subiculum (D). (E) Example images from *App^NL-G-F^Cst7^+/+^* and App^NL-G-F^*Cst7*^−/−^ subicula in male and female mice. Top panel shows LAMP2 (cyan) and bottom panel shows IBA1 (green), 6E10 (magenta) and LAMP2 (cyan) merge. Scale bars are 50 μm. (F-H) Ratio of IBA1/LAMP2 double positive staining *vs.* IBA1 total staining in the cortex (F), hippocampus (G), and dorsal subiculum (H) of wild-type, *App^NL-G-F^Cst7^+/+^* and *App^NL-G-F^Cst7*^−/−^ mice. (I&J) Example images from the cortex (I) and subiculum (J) of an *App^NL-G-F^* mouse stained with 6E10 (magenta) and MeX04 (white). (K) Ratio of MeX04 total staining *vs.* 6E10 total staining in cortex (blue), hippocampus (red) and subiculum (green) of male and female *App^NL-G-F^* mice. Scale bars are 50 μm. Data points are average from 2 fields of view/mouse (1 for subiculum) and bars are plotted as mean + S.E.M. n=5-12. *p<0.05, **p<0.01, ***p<0.001 by 2-way ANOVA with Tukey’s multiple comparisons post-hoc test.

To investigate the effect of *Cst7* knockout on lysosomal activity, we quantified LAMP2 immunostaining and normalized to IBA1 as these markers are highly correlated (Fig. S3E) and we wanted to remove the confounding factor of a greater IBA1 burden in *Cst7* knockout mice (Fig. 3B-D). Strikingly, we found in the subiculum that microglial-normalised LAMP2 staining followed the same pattern as many of the genes identified by microglia RNASeq with reduced expression in *App^NL-G-F^Cst7^−/−^ vs. App^NL-G-F^Cst7^+/+^* males but increased expression in *App^NL-G-F^Cst7^−/−^ vs. App^NL-G-F^Cst7^+/+^* females (Fig. 3E, quantification in Fig. 3H). A similar pattern was observed in the cortex and hippocampus but without a significant change in females (Fig. 3F&G). We hypothesized that the reason for this region-dependent effect may be due to differences in plaque structure. Aβ plaques typically comprise of a β-sheet-containing dense core and a more diffuse outer ring (DeTure and Dickson, 2019). We utilized Congo Red-derivative methoxy-X04 (MeX04), which binds selectively to fibrillar β-sheet-rich structures such as the dense cores of amyloid plaques (DeTure and Dickson, 2019), compared to anti-Aβ antibody 6E10, which will bind both dense cores and diffuse outers to study plaque composition in different brain regions of *App^NL-G-F^* mice (Fig. S3A). MeX04 was injected intraperitoneally 2.5 h before mice were sacrificed and brains taken for histology. We found that Aβ plaques had a far greater 6E10:MeX04 ratio in the subiculum compared to other brain regions (Fig. 3H&I, quantified in Fig. 3J), most likely caused by earlier Aβ accumulation in this region (Rönnbäck *et al.*, 2011; Gail Canter *et al.*, 2019) that may be dependent on a subpopulation of microglia (Spangenberg *et al.*, 2019). This implies that the sex-dependent effects of *Cst7* knockout on microglial lysosomes may be driven by plaque composition/structure, with regions dominated by dense core plaques (indicating further ‘disease progression’) showing greater effect of *Cst7* knockout.

### Cst7 regulates microglial Aβ burden in female App^NL-G-F^ mice

After discovering that *Cst7* knockout leads to sex-dependent changes in microglia at both the transcriptome and protein level indicative of an altered endolysosomal system, we next further investigated potential functional changes in these altered microglia. A major function of microglia in the context of AD is uptake of Aβ into lysosomes (Grubman *et al.*, 2021). To study this, we utilised MeX04 injection as described previously to label Aβ-containing microglia in the brain. First, we validated that intraperitoneal injection of MeX04 2.5 h before microglial isolation by FACS can detect Aβ-containing microglia. Wild-type and *App^NL-G-F^* mice were aged to 12 months and injected with either MeX04 or vehicle before microglial isolation by FACS as previously but with additional detection of MeX04 in the DAPI channel. As expected, wild-type + MeX04 and *App^NL-G-F^* + vehicle mice displayed a background MeX04 signal that was shifted only in the *App^NL-G-F^* + MeX04 group (Fig. S4).

After validating our approach, we next investigated the role of *Cst7* on microglial Aβ burden (Fig. 4A). Consistent with our RNASeq and histology data suggesting increased lysosomal capacity specifically in female *App^NL-G-F^* mice following deletion of *Cst7*, we observed a greater Aβ burden in female *App^NL-G-F^Cst7^−/−^* microglia compared to *App^NL-G-F^Cst7^+/+^* counterparts measured both by % Aβ-positive microglia (Fig. 4B) and mean fluorescence intensity of MeX04 on microglia (Fig. 4C). Interestingly, despite a reduced expression of lysosomal membrane marker LAMP2 in male *App^NL-G-F^Cst7^−/−^* mice compared to *App^NL-G-F^Cst7^+/+^* counterparts (Fig. 3C&D), we observed no difference in microglial Aβ burden (Fig. 4B). This may be due to the counteracting effect of a reduction of inflammatory genes such as *Il1b* and *Nlrp3* (Fig. 2G) which have been shown to impair phagocytosis (Heneka *et al.*, 2013; Tejera *et al.*, 2019).

**Figure 4.**
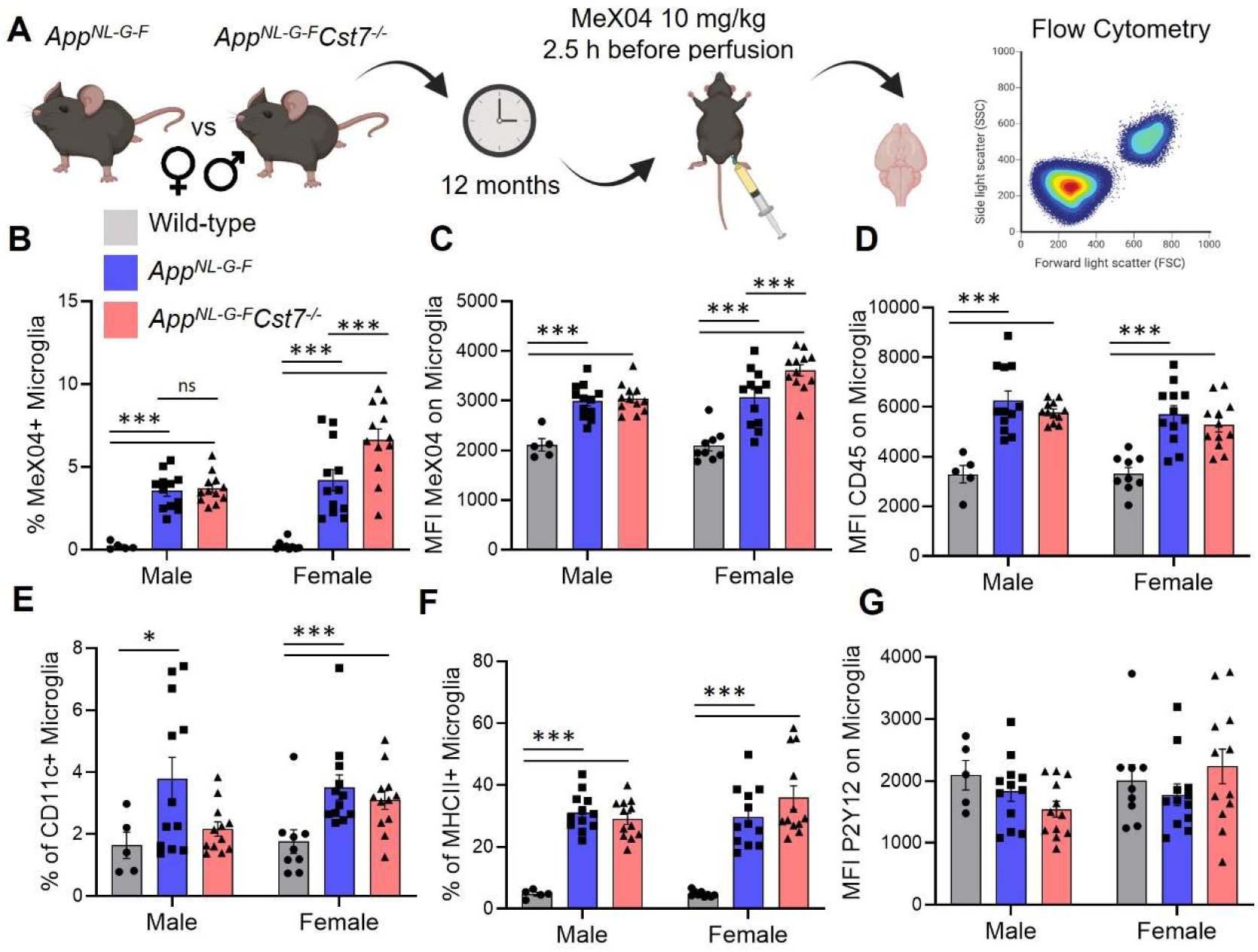
*Cst7* regulates microglial Aβ burden in female *App^NL-G-F^* mice. (A) Study design schematic. (B) % MeX04+ microglia of total microglia in male and female wild-type (grey), *App^NL-G-F^Cst7^+/+^* (blue) and *App^NL-G-F^Cst7*^−/−^ (red) brains. (C) Median fluorescence intensity (MFI) of MeX04 in microglia. (D) MFI of CD45 on microglia. (E) % CD11c+ microglia of total microglia. (F) % MHC-II+ microglia of total microglia. (G) MFI of P2Y12 on microglia. Bars are plotted as mean + S.E.M. n=5-12. *p<0.05, **p<0.01, ***p<0.001 by 2-way ANOVA with Tukey’s multiple comparisons post-hoc test.

The benefit of using a FACS-based approach to investigating microglial Aβ burden is the opportunity to include surface phenotypic markers to assess microglial reactivity. We assessed expression of CD45, a crude marker of overall microglial activation intensity; P2Y12, a microglial homeostatic marker; CD11c, a marker shown to be upregulated in disease/damage-associated microglia (Keren-Shaul *et al.*, 2017); and MHCII, a marker of antigen-presentation, a process which is also dependent on peptide processing through the endolysosomal system (Fig. S1). *App^NL-G-F^* microglia had increased expression of CD45, CD11c and MHCII compared to wild-type counterparts (Fig. 4D-F). We did not detect a reduction in P2Y12 expression (Fig. 4G). *Cst7* knockout did not lead to any significant differences in detection of these markers, although we did note a trend towards increase MHCII+ microglia only in female mice, perhaps consistent with elevated endolysosomal activity and our RNASeq data suggesting an increase in antigen-presentation pathway genes specifically in this group.

### Cst7 negatively regulates phagocytosis in microglia from female App^NL-G-F^ mice

Our current data suggest that *Cst7* plays a sex-dependent regulatory role on microglia leading to increased endolysosomal gene pathways, region-dependent lysosomal activity, and Aβ burden in female *App^NL-G-F^Cst7^−/−^* microglia. However, *in vivo* it is challenging to ascertain whether the increase in microglial Aβ is due to an increased uptake or to an impaired degradation. To test this, we conducted *in vitro* assays on *Cst7^+/+^* and *Cst7^−/−^* microglia to examine phagolysosomal and related functions. Initially, we examined microglia from young adult *Cst7^+/+^* and *Cst7^−/−^* background-matched mice to understand any baseline differences in the absence of a disease exposure. Unlike constitutively highly expressed microglial genes such as *Trem2* where knockout leads to dramatic alterations in *in vitro* phenotype (Takahashi, Rochford and Neumann, 2005; Hsieh *et al.*, 2009; Kleinberger *et al.*, 2014; Yeh *et al.*, 2016), *Cst7* is expressed at very low levels basally ((Sala Frigerio *et al.*, 2019)/Fig. 1E). Therefore, we reasoned that *Cst7* deletion would have limited impact on microglial phenotype at baseline where *Cst7* is not highly expressed. Comparing *Cst7^+/+^* and *Cst7^−/−^* microglia, we observed no significant difference in uptake of Aβ oligomers, human AD synaptoneurosomes, *S.Aureus* bioparticles, or myelin debris (Fig. S5A-D); degradation of Aβ (Fig. S5E); or lysosomal hydrolysis measured by the ability to cleave the quencher molecule from DQ-BSA (Fig. S5F); transcriptional response to LPS, IL-4 or apoptotic neurons (Fig. S5G-I); cytokine secretion in response to LPS or IL-4 (Fig. S6) or NLRP3 inflammasome activation by silica particles (Fig. S5J&K). We also addressed the possibility of compensation in adult knockout mice by using siRNA to knock-down *Cst7* in the BV2 microglia cell line and found that, despite 70% reduction in gene expression (Fig. S7A), *Cst7* knockdown had no effect on IL-6 secretion (Fig. S7B) or transcriptional changes response to LPS or IL-4 (Fig. S7C-E).

Therefore, in order to investigate CF function *in vitro* in the context of the disease-associated marked induction of microglial *Cst7* expression (see above) we developed a model system using primary microglia isolated from 12-month *App^NL-G-F^* mice and cultured them for 3 days *in vitro* (Fig. 5A). We found that microglia isolated from *App^NL-G-F^* mice then cultured for 3 days express 20-fold more *Cst7* than from age-matched wild-type counterparts (Fig. 5B). We also found that these *App^NL-G-F^* microglia express 2-fold less *Tmem119*, an established microglial homeostatic gene, than wild-type cells (Fig. 5C). These data suggest that, despite removal from a disease brain and subsequent culture *in vitro*, 12-month-old *App^NL-G-F^* microglia retain key elements of their *in vivo* disease transcriptional identity.

**Figure 5.**
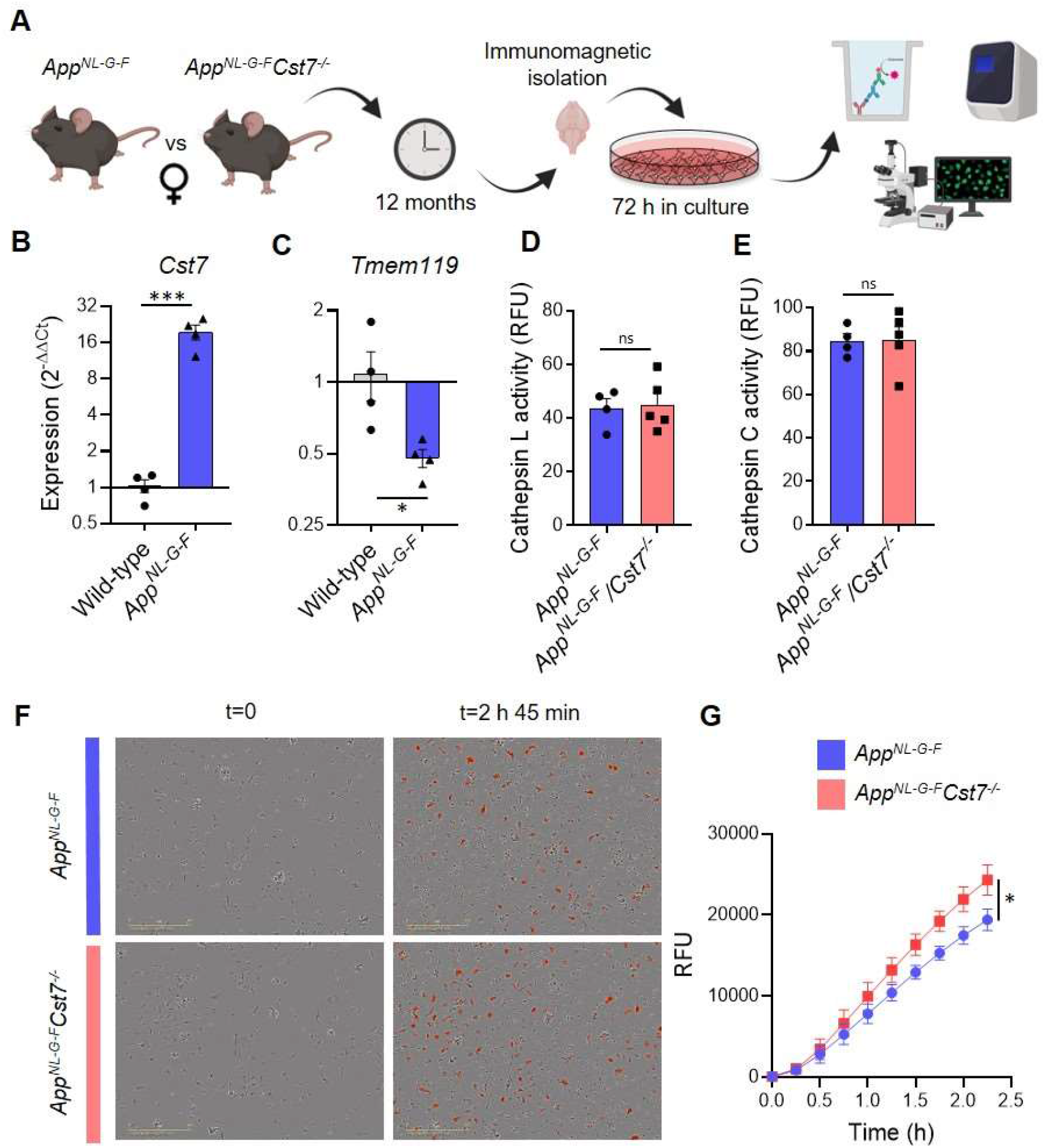
*Cst7* negatively regulates phagocytosis in microglia from female *App^NL-G-F^* mice. (A) Study design schematic. (B-C) qPCR data for *Cst7* (B) and *Tmem119* (C) RNA expression from female wild-type and *App^NL-G-F^* microglia isolated by CD11b beads and cultured for 3 days *in vitro*. Bars represent mean fold-change calculated by ΔΔCt. n=4. *p<0.05, ***p<0.001 by two-way t-test. (D-E) Cathepsin L (D) and C (E) activity from female *App^NL-G-F^Cst7^+/+^* (blue) and *App^NL-G-F^Cst7*^−/−^ (red) microglia measured by probe-based assay. Bars represent mean relative fluorescence units (RFU) + S.E.M. n=4-5. (F) Example images from myelin phagocytosis assays with female *App^NL-G-F^Cst7^+/+^* (blue) and *App^NL-G-F^Cst7*^−/−^ (red) microglia taken immediately before (t=0) and 2 h 45 min after (t=2 h 45 min) addition of pHrodo-tagged myelin. (G) Quantification of (F). Data shown are mean RFU (calculated as total integrated intensity of pHrodo Red normalised to cell confluence) +/− S.E.M. *p<0.05 calculated by unpaired t-test on area under curve (AUC) of data.

We then used this validated system to compare functional properties of microglia isolated from female *App^NL-G-F^Cst7^−/−^* and *App^NL-G-F^Cst7^+/+^* mice. First, we showed that *Cst7* knockout had no effect on isolation yield (Fig. S8A) and observed no differences in inflammatory cytokine secretion in response to LPS or NLRP3-activation stimulus silica (Fig. S8B-D), as predicted from our previous RNASeq analysis in female mice. Next, we tested whether *Cst7* plays a role in endolysosomal processes such as phagocytosis or lysosomal degradation. We used a pulse-chase live-imaging assay finding that *Cst7* knockout does not affect degradation of Aβ_1-42_ (Fig. S8E), a process dependent on cathepsins as it was inhibited by broad-spectrum inhibitors K777 and Ca074-Me (Fig. S8F). We also showed that *Cst7* deletion does not affect intracellular activity of cathepsins L and C (Fig. 5D&E), targets of cystatin-F (Hamilton *et al.*, 2008), or microglial lysosomal hydrolysis (Fig. S8G). These data would suggest that the increased microglial Aβ burden *in vivo* observed in Fig. 4A is not due to decreased degradative capacity and may be due increased phagocytosis. To test this, we performed uptake/phagocytosis assays on microglia isolated from 12-month female *App^NL-G-F^Cst7^−/−^ vs. App^NL-G-F^Cst7^+/+^*. To remove the confounding factor that female *App^NL-G-F^Cst7^−/−^* will already contain a greater more Aβ (Fig. 4B&C) and this may influence further Aβ uptake, we used myelin debris tagged with the pH-sensitive dye pHrodo red. We found that *Cst7* knockout led to a significant increase in microglial phagocytosis of myelin debris (Fig. 5F&G and Supplementary video 1). We also tested uptake with fluorescently tagged Aβ_1-42_ and found a trend towards increased uptake of Aβ_1-42_ (Fig. S8H).

Together, these data suggest that, in females, *Cst7/*CF negatively regulates phagocytosis but does not affect lysosomal proteolysis or PRR-driven inflammatory cytokine production in microglia from the *App^NL-G-F^* model of amyloid-driven AD.

### Effect of Cst7 deletion on disease pathology

Having discovered that *Cst7* plays sex-dependent regulatory roles on microglial phenotype and function in the *App^NL-G-F^* model of amyloid-driven AD, we sought to determine if this is associated with effects on disease pathology. Therefore, we studied the Aβ plaques in 12-month male and female *App^NL-G-F^Cst7^+/+^* and *App^NL-G-F^Cst7^−/−^* mice across brain regions. We observed the greatest plaque burden in the cortex and dorsal subiculum followed by the hippocampus (Fig. 6A) and, as expected, the cerebellum was largely devoid of plaque pathology. Next, we compared plaque burden within each area between *App^NL-G-F^Cst7^+/+^* and *App^NL-G-F^Cst7^−/−^* male and female mice. In males, there was no effect of *Cst7* genotype on plaque burden in any region (Fig. 6B-F). Surprisingly, despite an increased lysosomal burden (Fig. 3F) and intramicroglial Aβ load (Fig. 4B) in female *App^NL-G-F^Cst7^−/−^* mice, there was no decrease in amyloid plaque burden in any region. In fact, the reverse pattern was evident in some areas, most notably in the subiculum (Fig. 6D). In order to further probe the increase in amyloid pathology in the subiculum, we used MeX04-labelling of the dense-cores of Aβ plaques and counted the number of plaques in this region. We observed a significant although modest increase in plaque number specifically in the females (Fig. 6G, quantification in H) without any difference in average plaque size detected (Fig. S9). Together, these data suggest that *Cst7*/CF plays a negative regulatory role on microglial endolysosomal function in female AD mice and show that removal of this block and resultant increase in phagocytosis in *App^NL-G-F^Cst7^−/−^* females is associated with increased plaque pathology.

**Figure 6.**
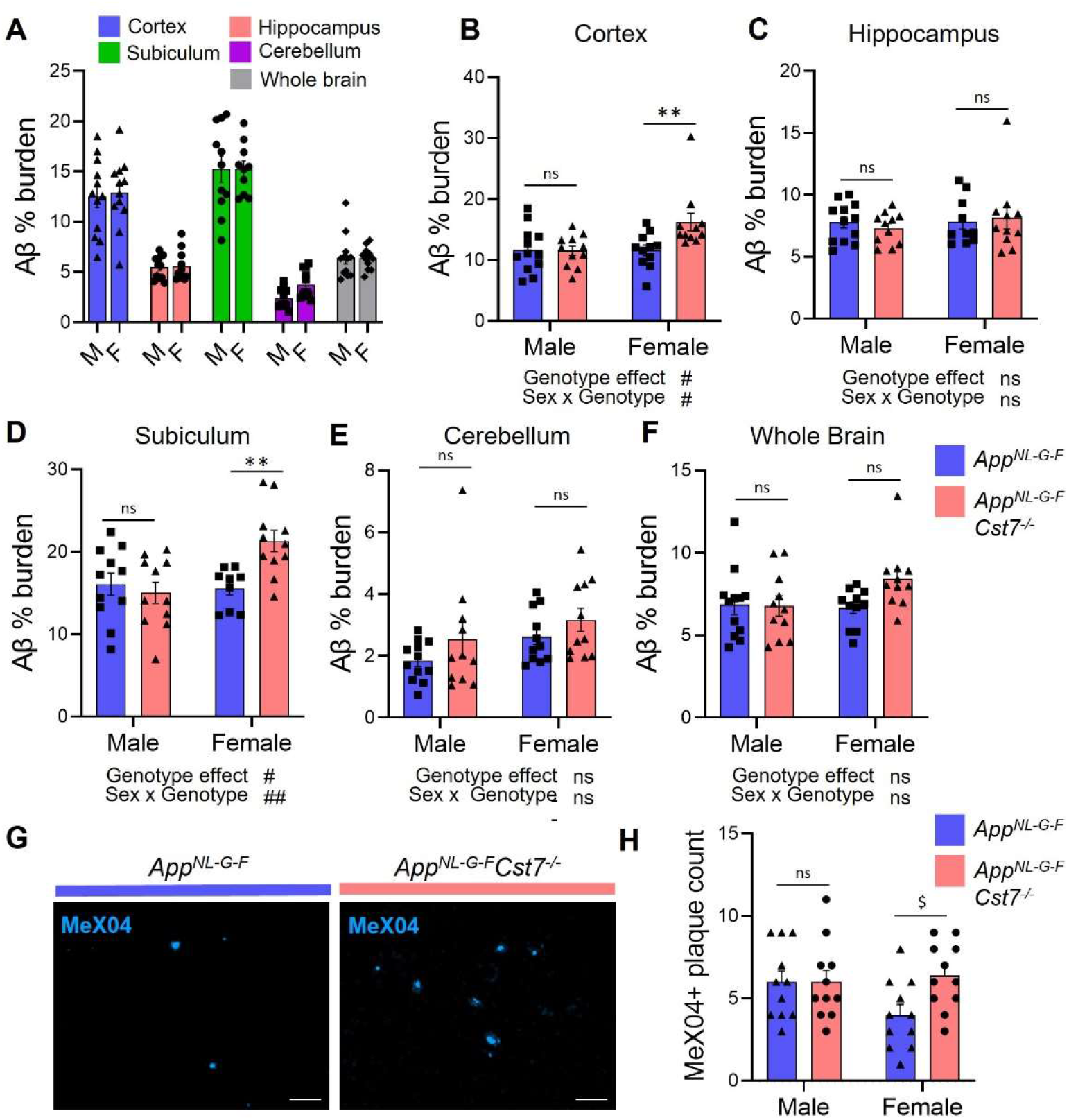
Investigating the effect of Cst7 deletion on plaque pathology. (A) % Aβ coverage measured by 6E10 DAB-staining in cortex (blue), hippocampus (red), subiculum (green), cerebellum (magenta) and whole brain (grey) of male and female *App^NL-G-F^* brains. (B-F) % Aβ coverage in the cortex (B), hippocampus (C), subiculum (D), cerebellum (E) and whole brain (F) of male and female *App^NL-G-F^Cst7^+/+^* (blue) and App^NL-G-F^*Cst7*^−/−^ (red) mice. Points are measured by thresholding of whole region area in sagittal section. Bars represent mean + S.E.M % coverage. n=10-12. **p<0.01 calculated by two-way ANOVA with Sidak’s multiple comparisons post-hoc test. #p<0.05, ##p<0.01 calculated by mixed effects analysis. (G) Example images from subicula of female *App^NL-G-F^Cst7^+/+^* (left) and *App^NL-G-F^Cst7*^−/−^ (right) mice stained with MeX04 (blue). Scale bars are 50 μm. (H) Quantification of (G). MeX04+ plaque count in the subicula of female *App^NL-G-F^Cst7^+/+^* (blue) and *App^NL-G-F^Cst7*^−/−^ (red) mice. Points are number of plaques from a single field of view for each mouse. Bars are mean plaque count + S.E.M. n-10-12.. $p<0.05 calculated by 2-way Mann-Whitney test.

## Discussion

Changes in microglial cells have been heavily implicated in the pathogenesis of AD, particularly supported by genetics studies, where expression of over 50% of risk genes is enriched in microglia (Hansen, Hanson and Sheng, 2018). Notably, a transcriptomic profile has emerged involving *Trem2*-dependent downregulation of homeostatic genes and upregulation of a signature characterised by genes involved in phagocytosis, lipid handling, and endolysosomal transport (Deczkowska *et al.*, 2018). One of the most robustly upregulated of these genes, *Cst7* (cystatin F), is believed to be involved in protease inhibition. However, the cellular functions controlled by *Cst7*/CF in microglia and whether Cst7/CF has a disease-modifying role in AD-related CNS proteinopathy, has not been explored previously. Here, we reveal that *Cst7*/CF plays a sexually dimorphic role in microglia in an amyloid-driven AD model, acting as a restraint on microglial endolysosomal activity and phagocytosis specifically in female mice, that when absent results in aggravated Aβ pathology.

Since their initial description in 2017, the DAM/MGnD/ARM gene signature has been extensively investigated with over 4 000 citations between the three papers describing them in five years. However, beyond gene ontology analysis, there is surprisingly little understanding of what many of the hallmark genes comprising this signature functionally do in the context of neurodegenerative disease. This is the first study to our knowledge to test the functional role of *Cst7*, one of the most consistently replicated markers of this cell state transcriptomic signature, in β-amyloid-driven pathology. Perhaps counterintuitively, *Cst7* (a reported cysteine protease inhibitor) is robustly upregulated in microglia alongside concomitant upregulation of cysteine proteases such as cathepsins D, B, L, and Z. In fact, such internal regulation systems are not uncommon in inflammation biology, where proinflammatory mediators such as IL-1β and IL-18 are negatively regulated by co-expressed IL-1 receptor antagonist and IL-18 binding protein, respectively (Hurme and Santtila, 1998; Dinarello *et al.*, 2013). We therefore hypothesise that *Cst7*/CF is upregulated along with numerous cathepsin genes in order to prevent potentially harmful protease activity (Turk *et al.*, 2012). In this study, we show that *Cst7*/CF knockout triggers an increase in microglial lysosomal activity and amyloid uptake. Intriguingly, this did not seem to depend on overactive intracellular cathepsin activity, which intuitively would have been expected to increase with the lack of negative regulator. It is possible that CF does in fact lead to an increase in cathepsin activity, but CF itself it is secreted and only impairs secreted cathepsins which are responsible for promoting phagocytosis. Indeed, studies have suggested that CF can be secreted and that there is a role for secreted cathepsins in phagocytosis (Liuzzo, Petanceska and Devi, 1999; Hamilton *et al.*, 2008).

Another key implication from this study is that of sexual dimorphism in disease. Here, we show stark differences in the effect of *Cst7* deletion between males and females at both the gene and protein level. Although there is a relative paucity of studies investigating male *vs.* female microglia in homeostasis and disease, numerous differences have been identified (Hanamsagar and Bilbo, 2016; Lynch, 2022). In general, male microglia are believed to have higher baseline activity in processes such as inflammation, antigen presentation, and phagocytosis; whereas female microglia are more associated with neuroprotection and the DAM/ARM/MGnD signature in ageing and disease (Guneykaya *et al.*, 2018; Kodama and Gan, 2019; Sala Frigerio *et al.*, 2019; Villa, Della Torre and Maggi, 2019; Guillot-Sestier *et al.*, 2021). Female microglia also develop ‘faster’ than male microglia resulting in differential response to inflammatory stimuli (Hanamsagar *et al.*, 2017). These differences are only partially dependent on hormonal signalling as masculinisation of females by ovariectomy and hormone replacement only partially recapitulated male microglia (Villa *et al.*, 2018). Indeed, sexual dimorphism of microglia in adults appears a cell-intrinsic phenomenon as female microglia transplanted in male brains retain their transcriptional profile and reduce infarct size in males after ischemic stroke (Villa *et al.*, 2018). Interestingly, we generally do not observe marked sexual dimorphism in microglia in wild-type or *App^NL-G-F^* mice. However, DAM/ARM/MGnD gene expression was higher in female *vs.* male microglia and lysosomal activity was markedly higher in male *vs.* female brains measured by LAMP2 staining, which is consistent with the literature (Guneykaya *et al.*, 2018; Sala Frigerio *et al.*, 2019). Importantly, we observed a substantial interaction effect of *Cst7* deletion and sex, whereby removal of the *Cst7* gene led to an ‘unlocking’ of sexual dimorphism in our cohort. This is most clearly demonstrated by differentially expressed genes between male *vs.* female microglia rising from 33 in *App^NL-G-F^Cst7^+/+^* mice to 240 in *App^NL-G-F^Cst7*^−/−^ mice. This sex-dependent interaction effect has previously been observed in studies investigating the cystatin/cathepsin system, where deletion of *Cst3* (cystatin C) led to protection in the EAE-model of multiple sclerosis in females but not males (Hoghooghi *et al.*, 2020).This effect appeared to be sensitive to hormones as ovariectomy or castration followed by testosterone or estrogen/progesterone administration effectively reversed the effect of *Cst3* knockout. Indeed, sex:genetics interactions are not uncommon in neurodegenerative disease, with various risk variants posing greater or lesser effect on risk in men or women (Gamache, Yun and Chiba-Falek, 2020). One example of this interaction is *APOE*, in which the presence of the E4 allele confers a greater AD risk in women than in men (Altmann *et al.*, 2014).

### Ideas and Speculation

Here, we observed that *Cst7*/CF played a role in microglial amyloid uptake and endolysosomal gene expression only in females. Surprisingly, this did not lead to a decrease in Aβ plaque pathology as might be expected but an increase, although of modest degree, in specific brain areas. This discovery raises the important question of the relationship between microglial phagocytosis and plaques. It is generally considered that an increase in microglial phagocytosis should lead to reduction in plaque burden and benefit in disease (Heneka *et al.*, 2013; Tejera *et al.*, 2019). However, recent studies have emerged to suggest that phagocytosing microglia may in fact act to ‘build’ plaques rather than dismantle them. For example, microglia lacking TAM receptors Axl and Mer fail to take up Aβ which leads to an overall decrease in dense-core plaque formation (Huang *et al.*, 2021), microglia can ‘seed’ plaques from disease tissue into engrafted, non-affected regions (d’Errico *et al.*, 2022) and either genetic or pharmacological depletion of microglia leads to a reduction in plaque burden/intensity accompanied by increase in cerebral amyloid angiopathy (Spangenberg *et al.*, 2019; Kiani Shabestari *et al.*, 2022). Interestingly, only female mice were used in two of these studies (Huang *et al.*, 2021; d’Errico *et al.*, 2022), which parallels our sex-specific findings with *Cst7*. In the context of the above studies, where microglial depletion/inactivation leading to reduced parenchymal plaque burden is detrimental in disease, we speculate that *Cst7*/CF may contribute to restraining plaque formation by influencing microglial endolysosomal function.

### Limitations

Although these data provide novel insight into the sex-dependent role of microglial gene *Cst7* in AD, it is important to acknowledge some caveats of the study that point to potential future investigations into this complex biological phenomenon. Here, we investigated CF function solely in mouse. While human data on microglial CF is limited, some studies have identified CF expression in microglia around plaques in AD (Ofengeim *et al.*, 2017) and *CST7* transcript upregulation in AD-associated microglia clusters by single nucleus RNA sequencing (snRNASeq) (Gerrits *et al.*, 2021). However, CF/*CST7* has also remained undetected in similar immunostaining and snRNASeq studies (Nuvolone *et al.*, 2017; Zhou *et al.*, 2020). While the reason for this discrepancy is unclear, it may be partially due to sex. Indeed, there was a greater ratio samples from female brains in (Gerrits *et al.*, 2021) than in (Zhou *et al.*, 2020), as would be expected if *CST7* plays a more important role in female microglia. These data further demonstrate the importance of stratifying studies by sex. Additionally, while mechanistic studies investigating CF/*CST7* in human microglia differentiated from inducible pluripotent stem cells (iPSCs) would be valuable, our discovery that CF function is revealed in mouse microglia only when cells are taken from disease context would suggest more complex models are required. Finally, while this study details the discovery of the mechanistic role of CF/*Cst7* in an amyloid-driven AD model, the precise mechanism by which *Cst7* deletion affects microglia only in females, and the biological reason for the accelerated DAM/MGnD/ARM programme in females remains unknown. Future work should investigate this further with ovariectomy/hormone replacement and transplantation studies as those performed by (Villa *et al.*, 2018).

In summary, our data provide key mechanistic insight into the role of one of the most robustly upregulated genes in reactive microglia. These data suggest that *Cst7*/CF regulates some key aspects of microglial function, in a sex-dependent manner, and that these are associated with pathology-influencing effects in an amyloid-driven AD model that also manifest differently in males and females. We hypothesise that *Cst7*/CF plays a part in an internal regulatory system that balances the two crucial processes of phagocytosis and inflammatory signalling, which are sexually dimorphic processes. More broadly, and in view of the poorly understood functions of many other recently described hallmark microglial disease-related state mediators, it is imperative to consider interactions with sex when investigating their cellular and disease roles.

## Materials and Methods

### Animals

*App^NL-G-F^* mice were provided by the RIKEN BRC through the National BioResource Project of the MEXT/AMED, Japan. *Cst7^−/−^* mice were obtained from Colin Watts, University of Dundee, UK. C57Bl/6J mice used for myelin preparations were purchased from Charles River, UK. Animals were maintained under standard laboratory conditions: ambient temperatures of 21 °C (±2 °C), humidity of 40–50%, 12 h light/dark cycle, ad libitum access to water and standard rodent chow. Genotype groups were randomised during the study and experimenters were blinded to genotype during all experiments. All animal experiments were carried out in accordance with the United Kingdom Animals (Scientific Procedures) Act 1986 and approved by the Home Office and the local Animal Ethical Review Group, University of Edinburgh. Genotyping was performed using optimised assays from Transnetyx. Methoxy X-04 (MeX04, BioTechne) was reconstituted in DMSO at 10 mg/mL before diluting to 0.33 mg/mL in 6.67% Cremaphor EL (Fluka), 90% PBS (Merck). MeX04 was administered by intraperitoneal injection at 10 mg/kg 2.5 h before animals were terminated.

### Microglial FACS isolation

Microglia were isolated by enzymatic digestion and fluorescence-activated cell sorting as follows. Brains from 12-month-old male and female *App^Wt/Wt^Cst7^+/+^*, *App^NL-G-F^Cst7^+/+^* and *App^NL-G-F^Cst7^−/−^* mice were isolated by terminally anaesthetizing with 3% isoflurane (33.3% O_2_ and 66.6% N_2_O) and transcardial perfusion with ice-cold DEPC-treated 0.9% NaCl, 0.4% trisodium citrate. Brains were immediately separated down the midline then the right hemisphere placed into ice-cold HBSS (ThermoFisher) and minced using a 22A scalpel. Minced hemi-brains were then centrifuged (300 x g, 2 min) and digested using the MACS Neural Dissociation Kit (Miltenyi) according to manufacturer’s instructions. Briefly, brain tissue was incubated in enzyme P (50 μL/hemi-brain) diluted in buffer X (1900 μL/hemi-brain) for 15 min at 37 °C under gentle rotation before addition of enzyme A (10 μL/hemi-brain) in buffer Y (20 μL/hemi-brain) and further incubation for 20 min at 37 °C under gentle rotation. Following digestion, tissue was dissociated mechanically using a Dounce homogenizer (loose pestle, 20 passes) on ice and centrifuged (400 x g, 5 min at 4 °C). To remove myelin, tissue was resuspended in 35% isotonic Percoll (GE Healthcare) in HBSS overlaid with 1X HBSS and centrifuged (800 x g, 30 min, 4 °C, no brake). Following centrifugation, the supernatant and myelin layers were discarded and the pellet resuspended in 100 μL FACS buffer (PBS, 0.1% low endotoxin bovine serum albumin (BSA, Merck), 25 mM HEPES) and transferred to V-bottomed 96-well plates (ThermoFisher). Cell suspensions were treated with anti-mouse-CD16/32 to block Fc receptors (BioLegend, 5 μg/mL, 20 min at 4 °C) before washing and transferring to PBS and treating with Zombie NIR (BioLegend, 1:100, 15 min at RT) to label dead cells. Next, cells were stained with antibodies (Table 1) to identify microglia and incubated for 20 min at 4 °C. Antibodies (BioLegend) used were as follows:

**Table 1.** Antibodies used for flow cytometry.

| Fluorochrome | Antigen | Clone | Lot | Stock concentration (mg/mL) | Dilution |
| --- | --- | --- | --- | --- | --- |
| AlexaFluor488 | Ly6C | HK1.4 | B248739 | 0.5 | 500 |
| PE | P2Y12 | S16007D | B298459 | 0.2 | 200 |
| PEDazzle594 | MHCII | M5/114.15.2 | B216062 | 0.2 | 200 |
| PE-Cy7 | CD45 | 30-F11 | B271123 | 0.2 | 200 |
| APC | CD11c | N418 | B280313 | 0.2 | 200 |
| BV711 | CD11b | M1/70 | B305911 | 0.005 | 50 |

Following incubation, cells were washed with FACS buffer, resuspended in 500 μL FACS buffer and transferred to 5 mL round-bottom FACS tubes through cell strainer caps (BD). Cells were sorted through a 100 μm nozzle on a FACS Aria II (BD) at the QMRI Flow Cytometry Cell Sorting Facility, University of Edinburgh with the following strategy (Fig. S1). Debris were eliminated using forward scatter (area) *vs.* side scatter (area), singlets were isolated using forward scatter (area) *vs.* forward scatter (height), dead cells were eliminated with Zombie NIR, monocytes were eliminated by gating out Ly6C+ cells, and microglia were defined as CD11b+/CD45+. All antibodies were validated using appropriate fluorescence minus one (FMO) controls. Microglia were sorted directly into 500 μL RLT buffer (Qiagen) and 20 000 CD11b+/CD45+ events were collected for downstream analysis using FlowJo software.

### RNA purification, QC and sequencing

RNA was purified using RNEasy Plus Micro kits (Qiagen) according to manufacturer’s instructions. RNA was quantified and QC’d using a 4200 TapeStation System (Agilent) with a High Sensitivity RNA ScreenTape Assay according to manufacturer’s instructions. For *in vivo* experiments, RNA was pooled from 3-4 animals within the same experimental group. RNA within pools was equally distributed in amount between the animals. 1 ng of total RNA was taken forward for low-input RNA sequencing (Cambridge Genomics). Briefly, cDNA was generated using TakaraBio SMART-Seq v4 Ultra Low Input RNA kit before input into the Illumina Nextera XT library prep. Pooled libraries were sequenced on the NextSeq 500 (Illumina) using the 75 cycle High Output sequencing run kit, spiked with 5% PhiX at a depth of 30 million reads per sample.

### RNASeq analysis

Reads were mapped to the mouse primary genome assembly GRCm39, Ensembl release 104 (Cunningham *et al.*, 2022) using STAR version 2.7.9a (Dobin et al., 2013), and tables of per-gene read counts were summarised using featureCounts version 2.0.2 (Liao et al., 2014). Differential expression analysis was performed using DESeq2 (R package version 1.30.1) (Love et al., 2014), using an adjusted p-value cut-off of 0.05 to identify genes differentially expressed between conditions. Comparisons carried out were Male (M) *App^Wt/Wt^Cst7^+/+^ vs.* M *App^NL-G-F^Cst7^+/+^,* Female (F) *App^Wt/Wt^Cst7^+/+^ vs.* F *App^NL-G-F^Cst7^+/+^,* M *App^NL-G-F^Cst7^+/+^ vs.* M *App^NL-G-F^Cst7^−/−^*, F *App^NL-G-F^Cst7^+/+^ vs.* F *App^NL-G-F^Cst7^−/−^*, M *App^NL-G-F^Cst7^+/+^ vs.* F *App^NL-G-F^Cst7^+/+^*, M *App^NL-G-F^Cst7^−/−^ vs.* F *App^NL-G-F^Cst7^−/−^.* For each comparison Gene Ontology (GO) enrichment analysis was performed to provide insight into the biological pathways and processes affected; GO enrichment analysis was performed using topGO (Alexa, Rahnenführer and Lengauer, 2006)(R package version 2.42.0) Significantly up-regulated and down-regulated gene sets were compared against annotated GO terms from the categories of ‘biological process’. Selected statistically significant GO:BP terms were then visualised using GraphPad prism v9.

### qPCR

Cells were seeded at 75 000 cells/well in a 24-well plate 7 days before stimulation (primary microglia) or 250 000 cells/well in a 24-well plate 18 h before stimulation (BV2 cells). On the day of the assay, microglia were treated with vehicle (PBS, 24 h, Merck), LPS (100 ng/mL, 24 h, Merck), IL-4 (20 ng/mL, 24 h, R&D Systems) or apoptotic SH-SY5Y neuroblastoma cells (2:1 ratio neurons:microglia, 24 h) then lysed in RLT buffer and RNA isolated according to the manufacturer’s instructions. RNA (100-500 ng) was converted to cDNA using SuperScript™ IV Reverse Transcriptase (ThermoFisher) according to manufacturer’s instructions. qPCR was performed using PowerUp SYBR® Green PCR Master Mix (ThermoFisher) in 384-well format using an qTOWER^3^84 Real-time PCR machine (Analytik Jena). For BV2 cells, 10 ng cDNA (assuming 100% RNA to cDNA conversion) was loaded per well with 5 pmol primer/well in triplicate in a total volume of 10 μL. For primary microglia experiments, 0.5 ng cDNA was loaded per well. Data were normalized to the expression of the housekeeping gene *Gapdh* and were analysed using the ΔΔCt method. Primers used were as follows:

### Microglial bead-based isolation for *in vitro* studies

Primary adult mouse microglia were isolated and cultured as described previously (Grabert and McColl, 2018). Brains were isolated by terminally anaesthetizing with 3% isoflurane (33.3% O2 and 66.6% N2O) and transcardial perfusion with ice-cold 0.9% NaCl. Brains were immediately placed into ice-cold HBSS (ThermoFisher) and minced using a 22A scalpel before centrifugation (300 x g, 2 min) and digestion using the MACS Neural Dissociation Kit (Miltenyi) according to manufacturer’s instructions. Briefly, brain tissue was incubated in enzyme P (50 μL/brain) diluted in buffer X (1900 μL/brain) for 15 min at 37 °C under gentle rotation before addition of enzyme A (10 μL/brain) in buffer Y (20 μL/brain) and further incubation for 20 min at 37 °C under gentle rotation. Following digestion, tissue was dissociated mechanically using a Dounce homogenizer (loose pestle, 20 passes) on ice and centrifuged (400 x g, 5 min at 4 °C). To remove myelin, tissue was resuspended in 35% isotonic Percoll (GE Healthcare) overlaid with HBSS and centrifuged (800 x g, 40 min, 4 °C). Following centrifugation, the supernatant and myelin layers were discarded and the pellet resuspended in MACS buffer (PBS, 0.5% low endotoxin BSA (Merck), 2 mM EDTA, 90 μL/brain). Anti-CD11b microbeads (Miltneyi) were added (10 μL/brain) and the suspension incubated for 15 min at 4 °C before running through pre-rinsed (MACS buffer) LS columns attached to a magnet (Miltenyi). After washing with 12 mL MACS buffer, columns were removed from the magnet and cells retained (microglia) were flushed in 5 mL MACS buffer. Microglia were resuspended in Dulbecco’s Modified Eagle’s Medium/Nutrient Mixture F-12 (DMEM/F-12, ThermoFisher) supplemented with 100 U/mL penicillin and 100 μg/mL streptomycin (PenStrep, Merck), 10% heat-inactivated fetal bovine serum (FBS, ThermoFisher), 500 ng/mL rhTGFβ-1 (Miltenyi), 10 ng/μL mCSF1 (R&D Systems). Microglia were counted using a haemocytometer and plated out onto 24 or 96-well plates (Corning) coated with poly-L-lysine (Merck). Cells were cultured for 7 days with a half media change on day 3. For stimulation experiments, cells were stimulated with LPS, IL-4, or silica particles (US Silica) at concentrations and timepoints as described in each experiment in detail before supernatant was taken and assessed for IL-6, IL-1β or TNF-α by DuoSet ELISA (R&D Systems) as per manufacturer’s instructions.

### Immortalised cell culture

Murine BV2 microglial cells were cultured in Dulbecco’s Modified Eagle’s Medium (DMEM, ThermoFisher), 10% FBS (ThermoFisher) and 1% PenStrep (Merck). Human SH-SY5Y neuroblastoma cells were cultured in DMEM/F-12 (ThermoFisher), 10% FBS (ThermoFisher) and 1% PenStrep (Merck). To induce apoptosis, SH-SY5Y cells were plated in a 10 cm dish at 4 × 10^6^ cells/dish and stimulated with UV light (4 × 15 W UV bulb, 30 h) before incubation at 37 °C for 18 h. After incubation, cells were removed from dishes using Accutase® (Merck) and adjusted to 4 × 10^6^ cells/mL for stimulation onto microglia. All cells were cultured at 37 °C, 5% CO_2_ in a tissue culture incubator (Panasonic) and handled in a Safe 2020 Class II tissue culture cabinet (ThermoFisher).

### BV2 siRNA knockdown

Small interfering RNAs (siRNAs) (Silencer-Select, Ambion) targeting *Cst7* were used to induce gene knockdown in BV2 cells. Cells were plated out at 250 000 cells/well in a 24-well plate 18 h before knockdown. For transfection of each siRNA (*Cst7* and non-targeting control), a 1:1 ratio of Lipofectamine RNAiMAX (ThermoFisher) and the siRNA, both diluted in Opti-MEM (ThermoFisher), were added to desired cell supernatants, with 10 pmol siRNA and 1.5 μL Lipofectamine RNAiMAX used per well. After 24 hours, cells were stimulated with vehicle (PBS), LPS (100 ng/mL, 24 h, Merck) or IL-4 (20 ng/mL, 24 h, R&D Systems). Following stimulation, cells were lysed for RNA analysis and supernatants were analysed for IL-6 content by ELISA (DuoSet, R&D Systems) according to manufacturer’s instructions.

### Myelin purification and pHrodo tagging

Purified myelin was prepared from C57Bl/6J mouse brains as an adapted protocol from previously described (Erwig *et al.*, 2019). Briefly, brains were isolated, digested, and myelin layer fractionated using 35% percoll gradient as described above. Myelin was washed by dilution in HBSS and centrifuged at 400 x g for 5 min at 4 °C before suspending in 3 mL ice cold 0.32 M sucrose solution containing protease inhibitor cocktail (Roche). Next, myelin was layered onto 3 mL ice cold 0.85 M sucrose solution in a 10 mL ultracentrifuge tube (Beckman) before centrifugation at 75 000 x g for 30 min at 4 °C on a Beckman Ultracentrifuge with MLA-55 rotor (acceleration 7, deceleration 7). Myelin was collected from the 0.32 M: 0.85 M interface and placed in 5 mL ice cold ddH_2_O in a new Beckman ultracentrifuge tube and vortexed before further centrifugation at 75 000 x g for 15 min at 4 °C (acceleration 9, deceleration 9). Myelin was then reconstituted in 5 mL ddH_2_O and incubated on ice for 10 min before further centrifugation at 12 000 x g for 15 min at 4 °C (acceleration 9, deceleration 9). This wash and ddH_2_O incubation step was repeated before myelin was again fractionated in 0.32 M: 0.85 M sucrose gradient and washed as before. Finally, purified myelin was resuspended in 1 mL sterile PBS and stored at −80 °C until further use. Myelin was tagged with pHrodo Red according to manufacturer’s instructions. Briefly, myelin was centrifuged (10 000 x g 10 min) and resuspended in 100 μL pHrodo Red succinimidyl ester (100 μg/mL in PBS 1% DMSO). Myelin was incubated for 45 min at room temperature (RT) in the dark before washing x 2 with 1 mL PBS. Finally, tagged myelin was resuspended in 100 μL, aliquoted, and stored at −20 °C for further use.

### Microglial phagocytosis/degradation assays

Primary adult mouse microglia were isolated as described above and plated out in 96-well plates at 50 000 cells/well. Cells were cultured for 7 days or 3 days (in the case of cells from *App^NL-G-F^* brains) before stimulation. For phagocytosis assays, microglia were first imaged using an IncuCyte® S3 Live-Cell Analysis System (Sartorius) in ‘phase’ and ‘red’ channels to gain baseline information on confluence and background fluorescence. Next, cells were stimulated with Aβ_1-42_HiLyte647 (ThermoFisher, 0.5 μM), pHrodo Red *S. aureus* Bioparticles (ThermoFisher, 100 μg/mL), purified mouse myelin tagged with pHrodo Red succinimidyl ester (1:100, ThermoFisher) or isolated human synaptoneurosomes from AD brains (Tai *et al.*, 2014; Tzioras *et al.*, 2019) tagged with pHrodo Red succinimidyl ester (1:20, ThermoFisher) by media change. Immediately after stimulation, cells were returned to the IncuCyte S3 for imaging in the ‘phase’ and ‘red’ channel with 9 images/well every 15 min for 2-3 h. Images were analysed using the IncuCyte 2019B Rev2 software (Sartorius) as total integrated intensity in ‘red’ channel normalised to confluence in ‘phase’ channel.

### Human Tissue

Use of human tissue for synaptoneurosome experiments above was reviewed and approved by the Edinburgh Brain Bank ethics committee and the ACCORD medical research ethics committee, AMREC (approval number 15-HV-016; ACCORD is the Academic and Clinical Central Office for Research and Development, a joint office of the University of Edinburgh and NHS Lothian). The Edinburgh Brain Bank is a Medical Research Council funded facility with research ethics committee (REC) approval (11/ES/0022).

### Cathepsin probe assays

Cathepsin probe assays were carried out as described in (Hamilton *et al.*, 2008). Briefly, microglia were isolated and cultured for 3 days as above before cells were lysed in RIPA buffer (60 μL/well, Merck). Next, 25 μL lysate was combined with 175 μL cathepsin probe L (Z-Phe-Arg-AMC, 45 μM, Bachem) or C (H-Gly-Phe-AMC, 56 μM, Bachem) in an assay buffer comprising 150 mM NaCl, 2mM EDTA, 5 mM DTT and 100 mM trisodium citrate at pH 5.5 in a black-walled, black-bottom 96-well plate (Corning). Lysates (or lysis buffer alone as a background control) were incubated for 2 h at 37 °C before reading on a fluorescence plate reader (BMG LABTECH) at excitation 360 nm, emission 460 nm with gain adjusted as necessary. Data are presented as background-corrected as relative fluorescence units.

### Immunohistochemistry

Brains from 12-month-old male and female *App^Wt/Wt^Cst7^+/+^*, *App^NL-G-F^Cst7^+/+^* and *App^NL-G-F^Cst7^−/−^* mice were isolated by terminally anaesthetizing with 3% isoflurane (33.3% O_2_ and 66.6% N_2_O) and transcardial perfusion with ice-cold DEPC-treated 0.9% NaCl, 0.4% trisodium citrate. Brains were immediately separated down the midline then the left hemisphere placed into ice-cold 10% neutral buffered formalin (ThermoFisher). Hemi-brains were post-fixed for 48 h at 4 °C before processing for paraffin embedding. For fluorescence immunostaining used to investigate microglial and lysosomal burden, 6 μm sagittal sections were deparaffinised with 2 × 10 min xylene before rehydration in subsequent changes of 100%, 90% and 70% ethanol (all 5 min). Sections were rinsed in dH_2_O before antigen retrieval in 10 mM TrisEDTA pH 9 (30 min at 95 °C). Next, sections were rinsed and incubated with primary antibody (Table 3) in PBS, 1% BSA (Merck), 0.3% Triton-X (Merck) at 4 °C overnight. Sections were washed in PBS, 0.1% Tween-20 (Merck) 3 × 5 min before incubation with secondary antibodies (Table 3) in PBS, 0.1% Tween-20 (Merck), 1% BSA (Merck) at RT for 1 h. Sections were washed again, incubated in TrueBlack Lipofuscin Autofluorescence Quencher (1:300 in 70% ethanol, Biotium) for 1 min, washed again in PBS, and rinsed in dH_2_O before mounting with Fluorescence Mounting Medium (Dako). Slides were imaged using an AxioImager D2 microscope (Zeiss) with a 20X objective. For quantification, two random fields of view were taken per mouse for both cortex and hippocampus. One image was taken for subiculum. One mouse was excluded from analysis as the section taken did not contain hippocampus. To remove the influence of plaque burden on the results, images were taken of approximately equal 6E10 burden. Images were analysed using a threshold analysis with QuPath v0.3.0 (Bankhead *et al.*, 2017) and co-stained was quantified using Definiens Developer.

**Table 2.**
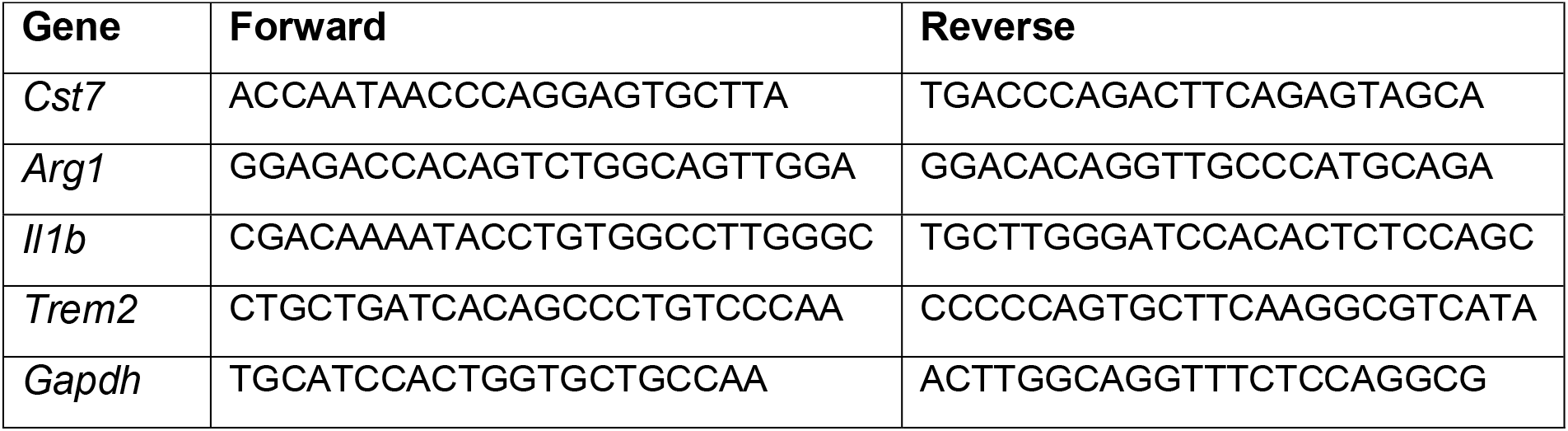
Primers used for qPCR.

**Table 3.** Antibodies used for immunohistochemistry.

| Antigen | Type | Species (raised) | Supplier | Stock concentration (mg/mL) | Dilution |
| --- | --- | --- | --- | --- | --- |
| IBA1 | Primary | Rabbit | Wako | 0.5 | 1:2000 |
| Aβ (6E10) | Primary | Mouse | BioLegend | 1 | 1:1000 |
| LAMP2 | Primary | Rat | BioLegend | 0.5 | 1:200 |
| Anti-rabbit IgG Alexa Fluor™ 488 | Secondary | Goat | ThermoFisher | 2 | 1:500 |
| Anti-mouse IgG<br>Alexa Fluor™<br>555 | Secondary | Donkey | ThermoFisher | 2 | 1:500 |
| Anti-rat IgG<br>Alexa Fluor™<br>647 | Secondary | Goat | ThermoFisher | 2 | 1:500 |
| Anti-mouse<br>Biotinylated | Secondary | Goat | Vector Labs | 1.5 | 1:100 |

For chromogenic 3, 3’-diaminobenzidine (DAB) staining used to quantify amyloid burden, 6 μm sagittal sections were deparaffinised and rehydrated as above. Following antigen retrieval, endogenous peroxidase activity was blocked with 0.3 % H_2_O_2_ for 10 min and sections were blocked with 5 % normal goat serum (Vector Labs) in PBS for 1 h at RT. 6E10 primary antibody was added at 1:500 and incubated overnight at 4 °C. The next day, sections were washed and stained with biotinylated anti-mouse IgG (Table 3) in PBST 1 % BSA for 1 h at RT before washing and addition of ABC Elite amplification (Vector Labs) for 30 min at RT according to manufacturer’s instructions. After a further wash, slides were immersed in DAB solution (0.5 mg/mL DAB, 0.015 % H_2_O_2_ in PBS) until stain had developed. Sections were counterstained with acidified Harris Hematoxylin (Epredia), dehydrated though increasing concentrations of ethanol and xylene, and mounted with Pertex mountant (CellPath). For imaging, slides were scanned with an Axioscanner Slide Scanner (Zeiss) in brightfield at 20X magnification. Images were analysed using a threshold analysis with QuPath v0.3.0 (Bankhead *et al.*, 2017). DAB slides were stained in two batches with equal representation of groups between batches, data are presented as batch-normalised 6E10 % positive area but analysis was carried out including batch as an independent variable in mixed effects analysis.

### Multiplex ELISA

Mouse cytokines from microglial supernatants were measured using MILLIPLEX® multiplex assays as described previously (McCulloch *et al.*, 2022). Briefly, MILLIPLEX MAP Mouse Cytokine/Chemokine Magnetic Bead Panel (MCYTOMAG-70K, Merck) was used to measure GM-CSF, IFN-γ, IL-1α, IL-1β, IL-2, IL-4, IL-5, IL-6, IL-12(p40), IL-33 and TNF-α. In all assays, samples were assayed as single replicates and all samples, standards and quality controls were prepared in accordance with the manufacturer’s instructions. Samples were incubated with beads on a plate for 1 h (isotyping assay) or overnight at 4 °C and washes carried out using a magnetic plate washer. Plates were analysed using a Magpix™ Luminex® machine and Luminex xPonent® software version 4.2, with a sample volume of 50 μL per well and a minimum of 50 events counted per sample.

### Randomisation and blinding

Experimenters were blinded to genotype groups throughout the study. Animals were given an experimental identifier and all samples were analysed using this coded identifier. Mice were randomly assigned to cull groups (which were performed in batches due to throughput for FACS) with stratification for experimental group. Order of MeX04 injection and subsequent mouse termination and tissue collection was randomised by random number generator using Microsoft Excel. Data was unblinded for analysis after experimental work was complete.

### Statistical analyses

Data are presented as mean values + standard error of the mean (S.E.M). Levels of significance were p<0.05 (*), p<0.01 (**), p<0.001 (***). Statistical analyses were carried out using GraphPad Prism (version 9). For RNASeq data, statistical significance was calculated using DESeq2 R package (Love, Huber and Anders, 2014). Immunohistochemistry, qPCR, cathepsin activity, and cytokine secretion were analysed with a two-way ANOVA followed by Tukey’s or Sidak’s post-hoc comparisons, unpaired Student’s t-test or mixed effects modelling. Live-imaging data were analysed by area under the curve followed by Student’s t-test or one-way ANOVA with Dunnett’s multiple comparisons test. Transformations were applied where necessary. Graphs and figures were created with GraphPad Prism (version 9), VolcaNoseR (Goedhart and Luijsterburg, 2020), and BioRender.com.

## Acknowledgments

This work was funded by the UK Dementia Research Institute which receives its funding from the Medical Research Council, Alzheimer’s Society, and Alzheimer’s Research UK. We gratefully acknowledge the contributions of our brain tissue donors and their families. BWM receives funding from the Medical Research Council (Grant Number MR/R001316/1) and from Leducq Foundation Transatlantic Network of Excellence, Stroke-IMPaCT (Grant Number 19CVD01). For the purpose of open access, the author has applied a CC-BY public copyright license to any Author Accepted Manuscript version arising from this submission. We would like to thank the QMRI Flow Cytometry & Cell Sorting Facility at The University of Edinburgh for assistance with FACS studies, the Shared University Research Facilities (SuRF) Histology facility at The University of Edinburgh for assistance with processing, sectioning and slide-scanning brains, Dr Daniel Soong at The MRC Centre for Reproductive Health, University of Edinburgh for his assistance with image analysis, and Prof. Siddharthan Chandran at The University of Edinburgh for providing the SHSY-5Y cell line.

## Author contributions

MJDD: designed experiments, performed and supervised experiments, analysed data, wrote the manuscript. LL: Managed *Cst7^−/−^* mouse colony, contributed intellectually to experiment planning. SS: Designed and optimised protocols for FACS isolation of microglia for RNASeq, isolated purified myelin. AD: Carried out siRNA knockdown experiments. LM: Carried out multiplex ELISA. JB: Contributed to analysis of published datasets. OD & XH: Analysed RNASeq data and contributed to analysis of published datasets. MM: Managed rodent colonies and genotyping. MT & TLS-J: Prepared synaptoneurosomes-enriched fractions from human AD brains and obtained/managed ethical approval for use of human tissue in the study. BM: Provided advice and evaluation of work, provided funding for work, aided in critical interpretation of work and edited drafts of the manuscript.

## Data availability

The data that support the findings of this study are available from the corresponding authors on request.

