## Supplementary Information for "Cystatin F *(Cst7)* drives sex-dependent changes in microglia in an amyloid-driven model of Alzheimer’s Disease"

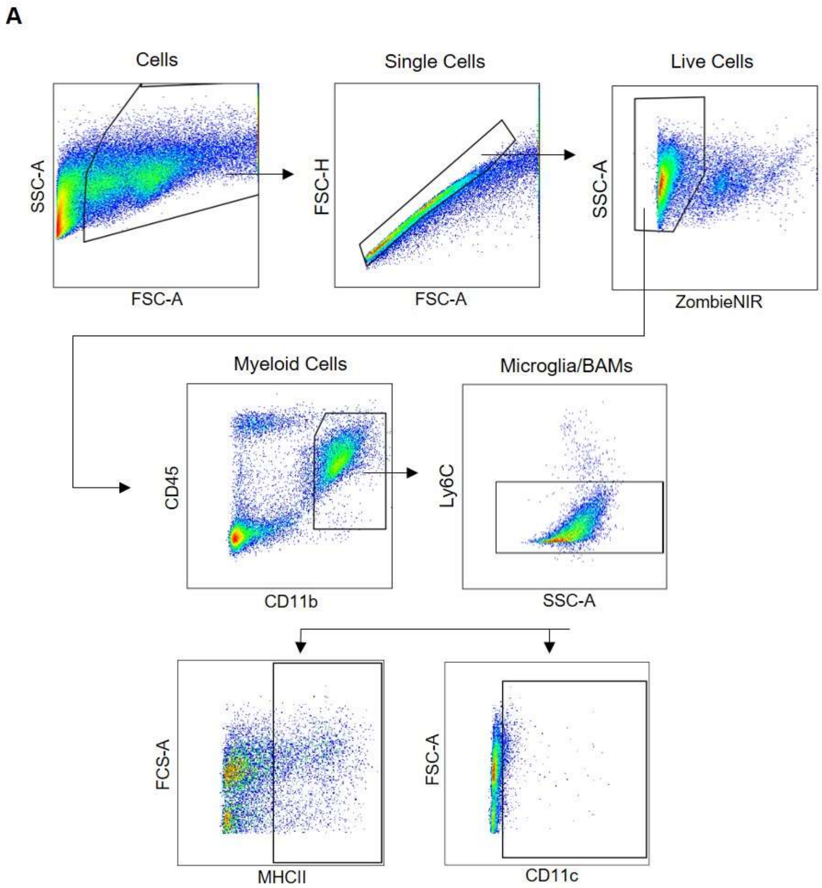


**Figure S1:** ***Gating strategy for microglia FACS.***

Microglia/BAMs were defined as single cells, Zombie NIR- live cells, CD11b+CD45+ myeloid cells, Ly6C- microglia/BAMs. Phenotypic markers MHCII and CD11c were also tested. All gates were set using fluorescence minus one (FMO) controls.

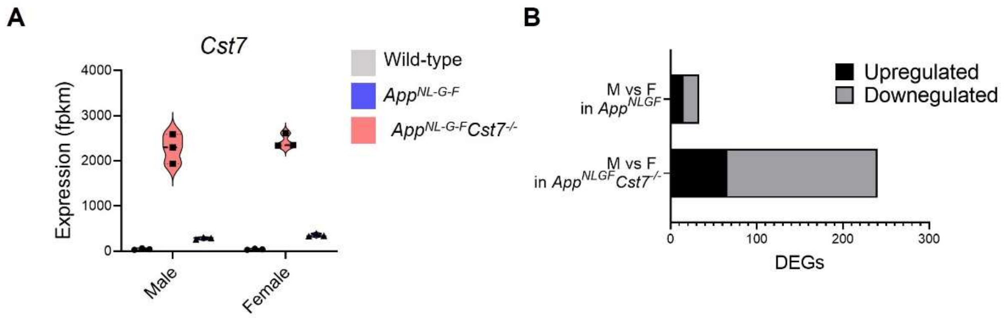


**Figure S2:** ***Gene expression from RNASeq data.***

(A) Example expression (fpkm) of *Cst7* from RNASeq of microglia isolated from male and female wild-type, $App^{NL\text{-}G\text{-}F}Cst7^{+/+}$ and $App^{NL\text{-}G\text{-}F}Cst7^{-/-}$ mice. (B) DEGs in male *vs.* female $App^{NL\text{-}G\text{-}F}$ or $App^{NL\text{-}G\text{-}F}Cst7^{-/-}$ mice.

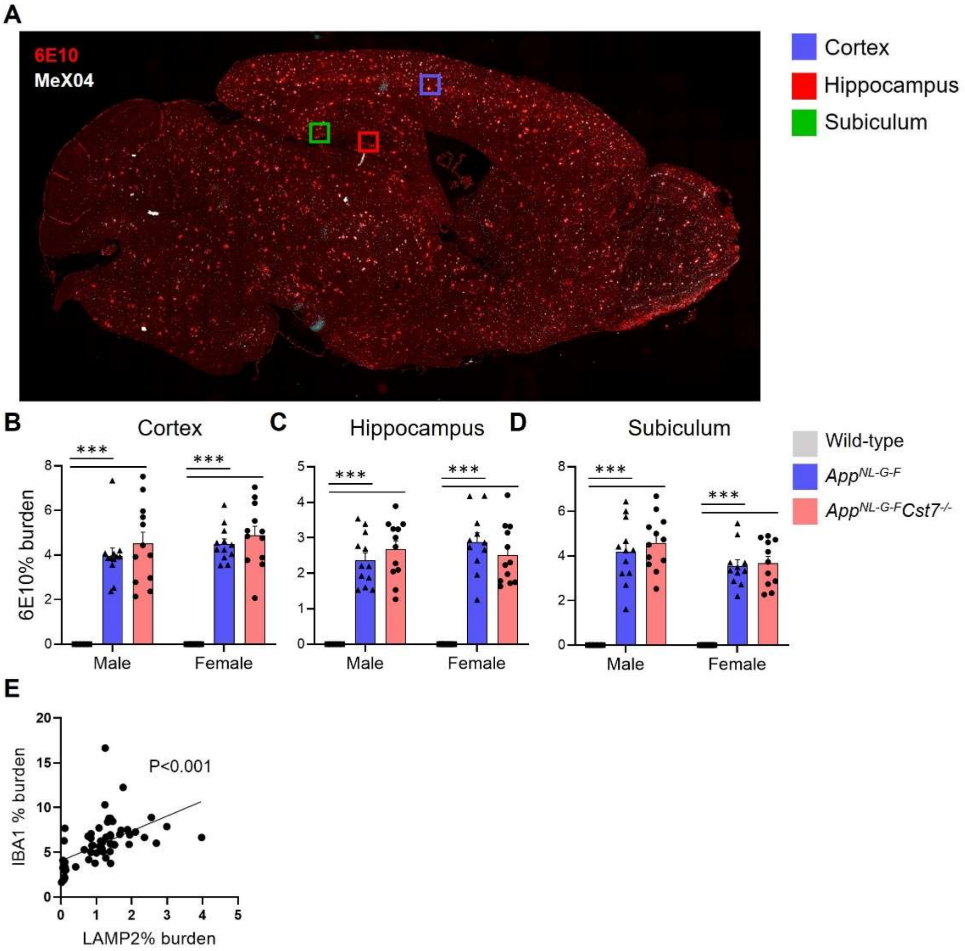


**Figure S3:** ***Immunohistochemistry supplementary information.***

(A) Whole slide-scan of an *App*$^{NL\text{-}G\text{-}F}$ brain stained with 6E10 (red) and MeX04 (white) to illustrate regions of interest taken for analysis. Example regions are cortex (blue square), hippocampus (red square) and subiculum (green square). (B-D) % 6E10 burden in cortex (B), hippocampus (C) and subiculum (D) of male and female wild-type, App$^{NL\text{-}G\text{-}F}$*Cst7*$^{+/+}$ and App$^{NL\text{-}G\text{-}F}$*Cst7*$^{-/-}$ mice. (E) % LAMP2 burden *vs.* % IBA1 burden scatter plot from all mice in study with linear regression. ***p<0.001 by 2-way ANOVA with Tukey's multiple comparisons post-hoc test.

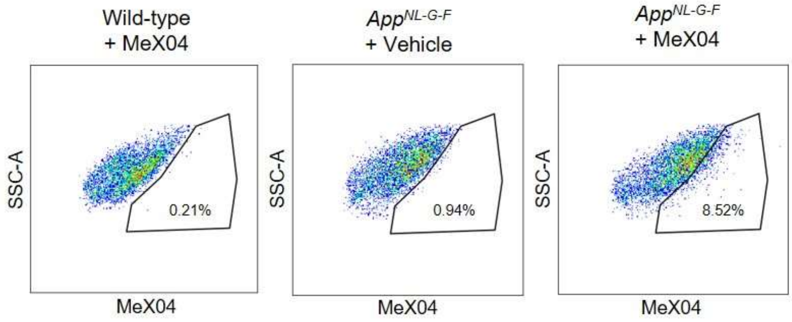


**Figure S4:** ***Validating MeX04 to measure Aβ-containing microglia***

FACS plots of microglia gated by MeX04 and SSC-A. Plots show data from a wild-type mouse injected with MeX04 (left), an aged App$^{NL-G-F}$ mouse injected with vehicle (middle), and an aged *App*$^{NL-G-F}$ mouse injected with MeX04 (right).

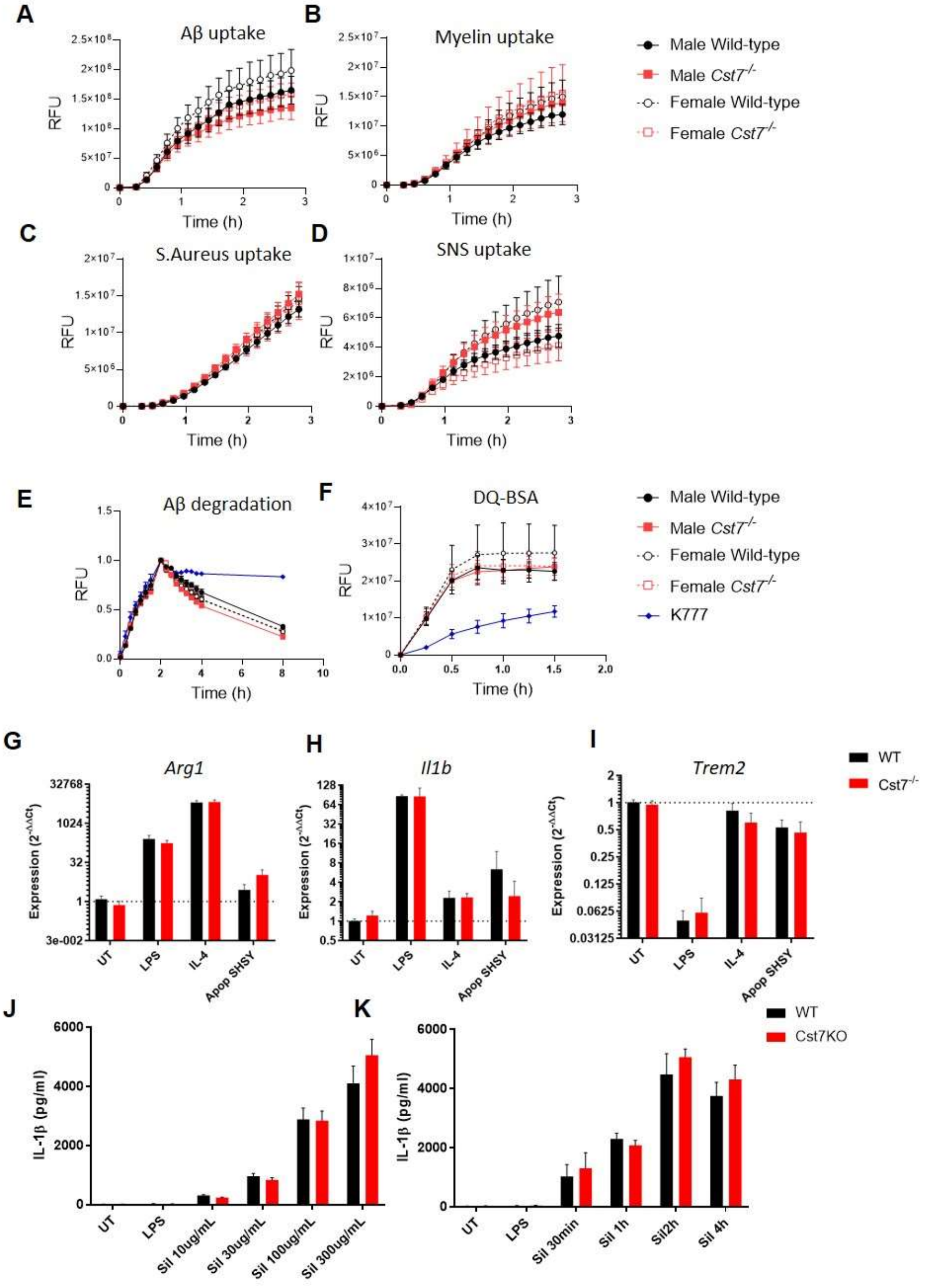


**Figure S5:** ***Investigating Cst7 knockout in vitro***

(A-D) Uptake of various tagged substrates from male (solid lines/symbols) and female (dotted lines/symbols) wild-type (black) and *Cst7*$^{-/-}$ (red) microglia. Substrates tested were HiLyte647-tagged A$\beta_{1-42}$(A), pHrodo-tagged myelin (B), pHrodo-tagged *S.Aureus* (C), and pHrodo-tagged human AD synaptoneurosomes (D). (E) Uptake and degradation of HiLyte647-tagged A$\beta_{1-42}$ from male (solid lines/symbols) and female (dotted lines/symbols) wild-type (black) and *Cst7*$^{-/-}$ (red) microglia and microglia treated with pan-cathepsin inhibitor K777 (10 µM, blue lines/symbols). (F) Lysosomal proteolysis measured by de-quenching of DQ-BSA from male (solid lines/symbols) and female (dotted lines/symbols) wild-type (black) and *Cst7*$^{-/-}$ (red) microglia and microglia treated with pan-cathepsin inhibitor K777 (10 μM, blue lines/symbols). Data are mean signal measured by RFU +/- S.E.M. n=4. (G-I) qPCR data showing gene expression (relative to wild-type untreated) of male wild-type (black) and *Cst7*$^{-/-}$ (red) microglia stimulated with LPS (100 ng/mL, 24 h), IL-4 (20 ng/mL, 24 h) or apoptotic SH-SH5Y cells (2:1 ratio, 24 h). Genes tested were *Arg1* (G), *Il1b* (H), and *Trem2* (I). (K&J) IL-1β secretion from male wild-type (black) and *Cst7*$^{-/-}$ (red) microglia treated with LPS (1 μg/mL, 4 h) or LPS + silica (varying dose, 4 h (K) or 300 μg/mL, varying time (J)) to activate the NLRP3 inflammasome. Data are mean secretion/expression + S.E.M. n=4

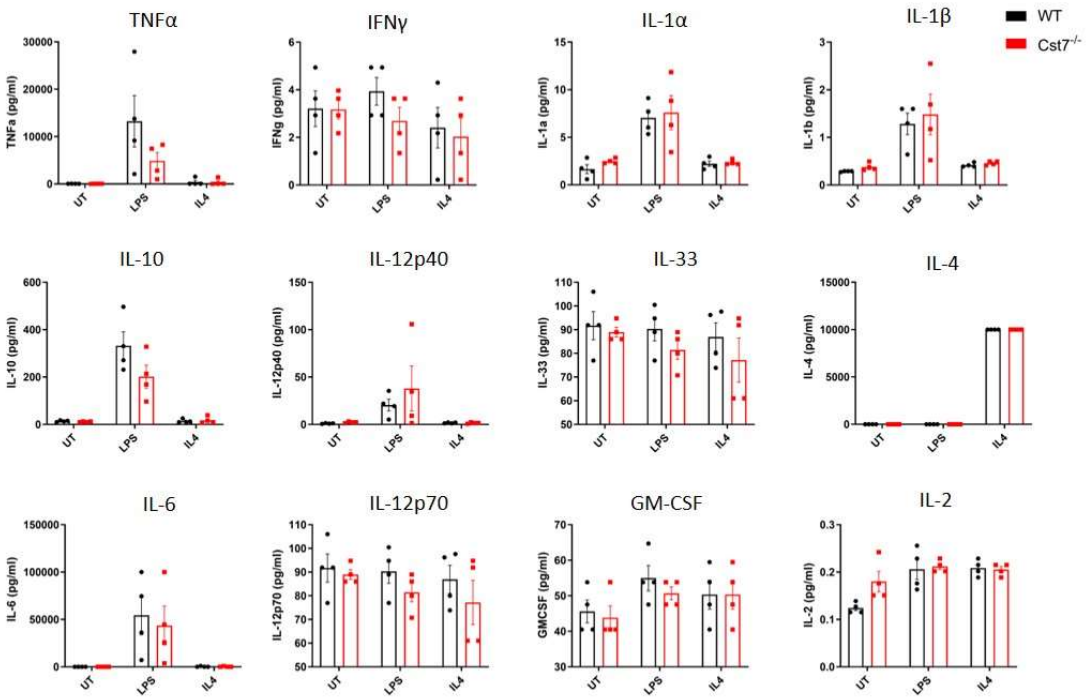


**Figure S6:** ***Investigating Cst7 knockout on microglial cytokine secretion***

Multiplex bead-based ELISA data from male wild-type (black) and *Cst7*$^{-/-}$ (red) microglia stimulated with LPS (100 ng/mL, 24 h) or IL-4 (20 ng/mL, 24 h). Data are mean +/- S.E.M. n=4.

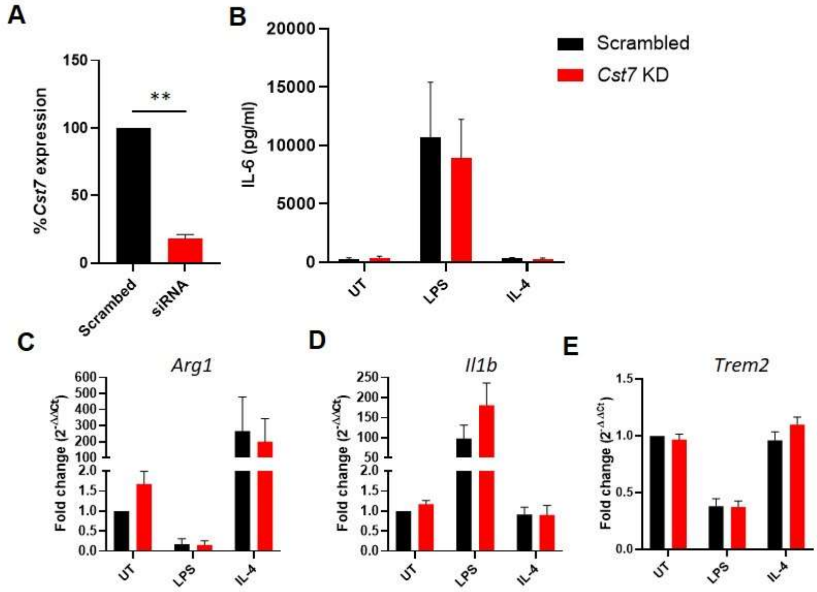


**Figure S7:** ***Investigating Cst7 knockdown in BV2 cells vitro***

(A) qPCR data of *Cst7* expression in BV2 cells treated with scrambled control siRNA (black) or siRNA targeting *Cst7* (red). (B) IL-6 secretion from BV2 cells treated with scrambled control siRNA (black) or siRNA targeting *Cst7* (red) following stimulation with LPS (100 ng/mL, 24 h) or IL-4 (20 ng/mL, 24 h). (C-E) qPCR data of gene expression in BV2 cells treated with scrambled control siRNA (black) or siRNA targeting *Cst7* (red) before stimulation with LPS (100 ng/mL, 24 h) or IL-4 (20 ng/mL, 24 h). Genes tested were Arg1 (C), Il1b (D) and Trem2 (E). **p<0.01 by one-sample t-test against hypothetical value of 100.

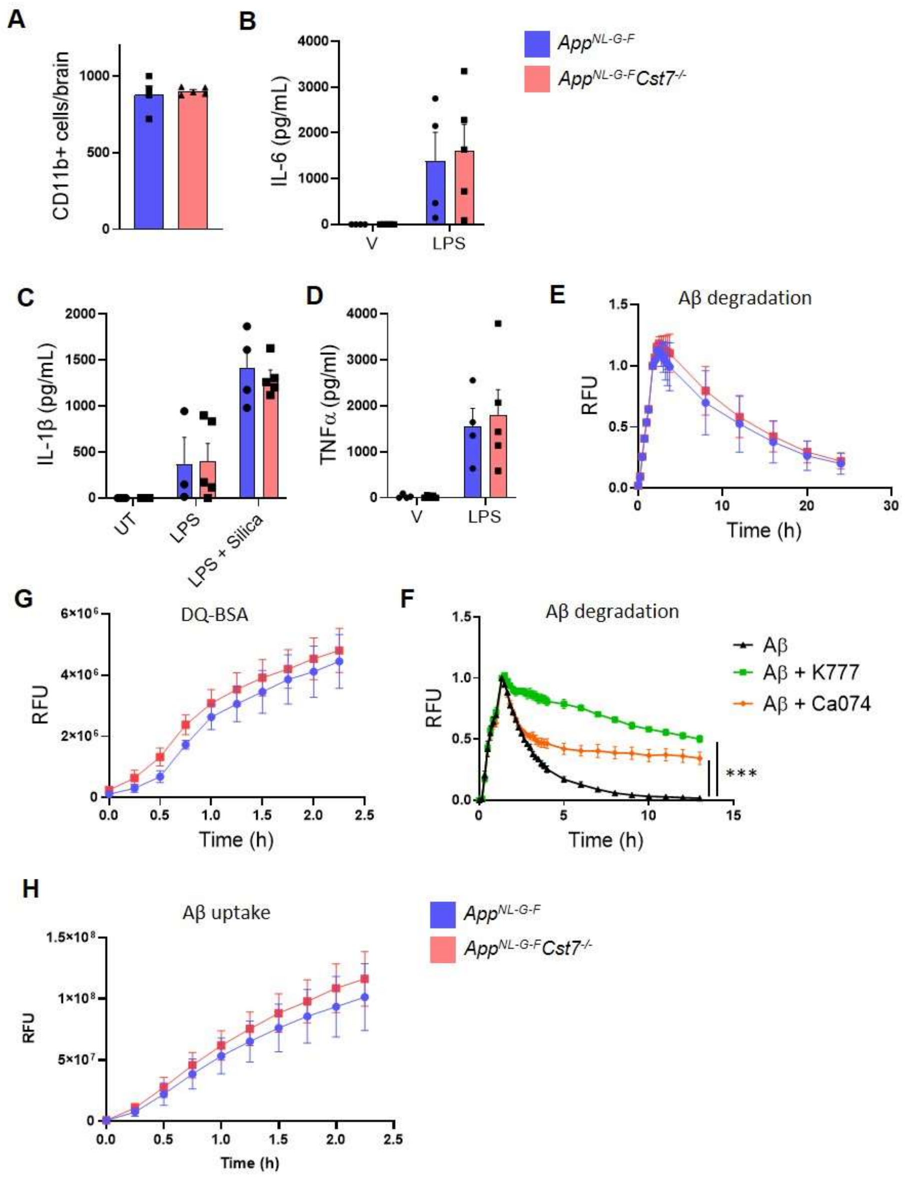


**Figure S8:** ***Investigating Cst7 knockout from App$^{NL\text{-}G\text{-}F}$ mice in vitro***

(A) Yield of isolated microglia from *App$^{NL\text{-}G\text{-}F}$Cst7$^{+/+}$* (blue) and App$^{NL\text{-}G\text{-}F}$*Cst7$^{-/-}$* (red) mice. (B) IL-6 secretion from *App$^{NL\text{-}G\text{-}F}$Cst7$^{+/+}$* (blue) and *App$^{NL\text{-}G\text{-}F}$Cst7$^{-/-}$* (red) cultured microglia treated with LPS (100 ng/mL, 24 h). (C) IL-1β secretion from *App$^{NL\text{-}G\text{-}F}$Cst7$^{+/+}$* (blue) and *App$^{NL\text{-}G\text{-}F}$Cst7$^{-/-}$* (red) cultured microglia treated with LPS (100 ng/mL, 24 h) or LPS (1 µg/mL, 4 h) + silica (300 µg/mL, 4 h) to activate the NLRP3 inflammasome. (D) TNFα secretion from *App$^{NL\text{-}G\text{-}F}$Cst7$^{+/+}$* (blue) and *App$^{NL\text{-}G\text{-}F}$Cst7$^{-/-}$* (red) cultured microglia treated with LPS (100 ng/mL, 24 h). Data are presented as mean cytokine concentration in pg/mL + S.E.M. (E) Uptake and degradation of HiLyte647-tagged A$β_{1\text{-}42}$ from female *App$^{NL\text{-}G\text{-}F}$Cst7$^{+/+}$* (blue) and *App$^{NL\text{-}G\text{-}F}$Cst7$^{-/-}$* (red) cultured microglia. (F) Uptake and degradation of HiLyte647-tagged A$β_{1\text{-}42}$ from BV2 cells treated with vehicle (0.5 % DMSO, black), K777 (10 µM, green) or Ca074Me (100 µM, orange). (G) Lysosomal proteolysis measured by de-quenching of DQ-BSA from female *App*$^{NL-G-F}$*Cst7*$^{+/+}$ (blue) and *App*$^{NL-G-F}$*Cst7*$^{-/-}$ (red) cultured microglia. (H) Uptake of HiLyte647-tagged A$\beta_{1-42}$ from female *App*$^{NL-G-F}$*Cst7*$^{+/+}$ (blue) and *App*$^{NL-G-F}$*Cst7*$^{-/-}$ (red) cultured microglia. Data are shown as mean +/- S.E.M. ***$p<0.001$ calculated by one-way ANOVA on area under curve (AUC) of data with Dunnett's multiple comparison's test.

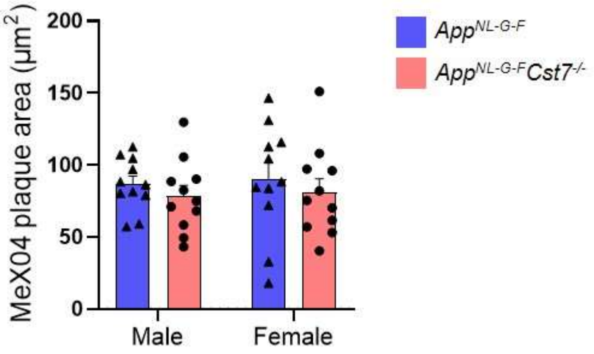


**Figure S9:** ***Investigating the role of Cst7 in plaque burden***

Area of MeX04+ plaques in the subiculum of male and female *App*$^{NL\text{-}G\text{-}F}$*Cst7*$^{+/+}$ (blue) and *App*$^{NL\text{-}G\text{-}F}$*Cst7*$^{-/-}$ (red) brains. Bars are mean plaque area + S.E.M.

**Table S1:** ***DEGs between male and female App$^{NL\text{-}G\text{-}F}$ and App$^{NL\text{-}G\text{-}F}$Cst7$^{-/-}$ mice (selected genes in bold)***

**A** Male vs Female in *App$^{NL\text{-}G\text{-}F}$*

| Up (higher in males) | Down (higher in females) |
|---|---|
| *Hykk* | *Gpx3* |
| *Nr1d2* | ***Gpnmb*** |
| *Ttc7* | *Gpx1* |
| *Gm14023* | *Rgs16* |
| *Pan3* | *Phlda3* |
| *Hyou1* | *Mgl2* |
| *Gm10382* | *Selenow* |
| *Tmem229a* | *N4bp3* |
| | ***Ctse*** |
| | *Kcnj2* |
| | *Lrrc27* |
| | ***Spp1*** |
| | *Gm33858* |
| | *Kcnk3* |
| | *Tmem51* |
| | ***C3*** |
| | *Cck* |

**B** Male vs Female in *App$^{NL\text{-}G\text{-}F}$Cst7$^{-/-}$*

| Up (higher in males) | | Down (higher in females) (selected) | | | |
|---|---|---|---|---|---|
| *Hsp90b1* | *Ccdc71* | *Mir155hg* | *Enpp2* | *Il12b* | *Col4a1* |
| *P4ha1* | *AI506816* | *Vegfa* | *Gpr37l1* | *Gadd45b* | *Rab20* |
| *Plod1* | *Mis12* | *Sox7* | *Armc2* | ***Tnf*** | *Csrnp1* |
| *Entpd1* | *Txndc5* | *Ttr* | *F3* | *Il1rn* | ***Nlrp3*** |
| *P2ry12* | *Stip1* | *Gdf3* | *Acsbg1* | *Gm1673* | *Dcstamp* |
| *Serinc3* | *Hsp90ab1* | *Gdf10* | ***Il1b*** | *Cd69* | *Lilrb4a* |
| *Selplg* | *Creld2* | *Id3* | *Hbegf* | *Lrguk* | *Cdkn1a* |
| *P2ry6* | *Rnf19a* | *Gpat2* | *Phlda1* | *Hes1* | *Tnfaip2* |
| *Pdia6* | *Pomt2* | ***Cxcl1*** | *Tamalin* | ***Nfkbid*** | *Nfkbiz* |
| *Tnfrsf11a* | *Exog* | ***Cxcl2*** | *Lzts3* | *Selenom* | *Insig1* |
| *P2ry13* | *Cog3* | *1200007C1* | *H2-K2* | *Sdc4* | *Pim3* |
| *Rpn1* | *Sec23ip* | *Nr4a1* | *Tmem178* | *Tmem51* | *Gem* |
| *Slc35e1* | *Tprn* | *Rgs11* | *Phlda3* | *Ccrl2* | *Gpx1* |
| *Pros1* | *Hyou1* | *Slc39a2* | *Bcl2a1d* | *Cd83* | *Miip* |
| *Eif2ak1* | *Zscan22* | *Ifrd1* | *Kcnh2* | ***Gpnmb*** | *Marcksl1* |
| *Fkbp4* | *Zfp516* | *H2-Q5* | *Maff* | *Tnfaip3* | *Selenoh* |
| *Tm9sf2* | *Katnb1* | *Dusp2* | *Snhg15* | ***Il1a*** | *Rel* |
| *Dusp6* | *Pigg* | *Mgl2* | *Gpx3* | *Gm13889* | *Vcam1* |
| *Pdia4* | | *Tnfsf9* | *Kdm6b* | *Osm* | ***Spp1*** |
| *Txndc11* | | *Selenow* | *Siglec1* | *Tob1* | *Id2* |
| | | *Dusp5* | *Cxcl10* | *Rasgef1b* | *Nfkbia* |
